# kCSD-python, reliable current source density estimation with quality control

**DOI:** 10.1101/708511

**Authors:** Chaitanya Chintaluri, Marta Bejtka, Władysław Średniawa, Michał Czerwiński, Jakub M. Dzik, Joanna Jędrzejewska-Szmek, Daniel K. Wójcik

## Abstract

Interpretation of the extracellular recordings can be difficult due to the long range of electric field but can be facilitated by estimating the density of current sources (CSD). Here we introduce `kCSD-python`, an open Python package implementing Kernel Current Source Density (kCSD) method, and introduce several new techniques to facilitate CSD analysis of experimental data and interpretation of the results. We investigate the limitations imposed by noise and assumptions in the method itself. `kCSD-python` allows CSD estimation for arbitrary distribution of electrodes in 1D, 2D, and 3D, assuming distributions of sources in tissue, a slice, or in a single cell, and includes a range of diagnostic aids. We demonstrate its features in a Jupyter notebook tutorial to facilitate uptake by the community.

## 1 Introduction

Extracellular potential recordings are a mainstay of neurophysiology. However, the long range of electric field still makes their interpretation challenging despite decades of our experience. Extracellular potential in tissue is produced by transmembrane currents. Its low-frequency part, called the Local Field Potential (LFP), is believed to mainly reflect dendritic processing of synaptic inputs (Nunez and Srinivasan, 2006; Buzsáki, Anastassiou, and Koch, 2012). To facilitate understanding of the processes underlying the recorded signal it is useful to estimate the density of transmembrane current sources (Current Source Density, CSD) (Pitts, 1952; Nicholson and Freeman, 1975; Mitzdorf, 1985; Einevoll et al., 2013; Gratiy et al., 2017). CSD gives direct access to the physiologically relevant information, which is often concealed in original data (Mitzdorf, 1985). The relation between the CSD and the extracellular potential can be described by the Poisson equation

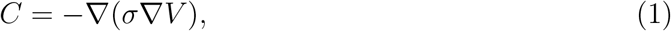

where *C* is the CSD, *V* is the extracellular potential, and *σ* — the conductivity tensor. For isotropic and homogeneous tissue Eq. (1) reduces to

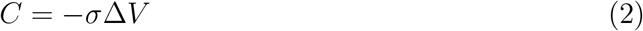

which can be solved:

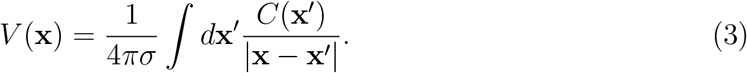

Eq. (2) and Eq. (3) show that knowing the potential in the whole space, we can compute the CSD, and knowing the CSD in the whole space, we can compute the potential. Experimentally, we can only access the potential at discrete electrode locations, so direct determination of the CSD in the whole space from Eq. (2) is impossible. To deal with this problem different methods for estimation of current sources have been proposed since the middle of the past century (Pitts, 1952; Nicholson and Freeman, 1975; Pettersen et al., 2006; Łęski, Wójcik, et al., 2007; Łęski, Pettersen, et al., 2011; Potworowski et al., 2012; Chintaluri et al., 2021; Klein et al., 2021). Here, we present `kCSD-python`, an open Python toolbox implementing the kernel Current Source Density (kCSD) method (Potworowski et al., 2012; Chintaluri et al., 2021). It allows kCSD reconstruction of current sources for data from 1D setups (laminar probes and equivalent electrode distributions), 2D (planar MEA, multishaft silicon probes, Neuropixel or SiNAPS probes, etc), and 3D electrode setups (Utah arrays, multiple electrodes placed independently in space with controlled positions), where the sources are assumed to come from tissue (kCSD) or from single cells with known morphology (skCSD, Cserpan et al., 2017). Fig. 1 shows the different experimental scenarios for which this software is applicable.

**Figure 1.**
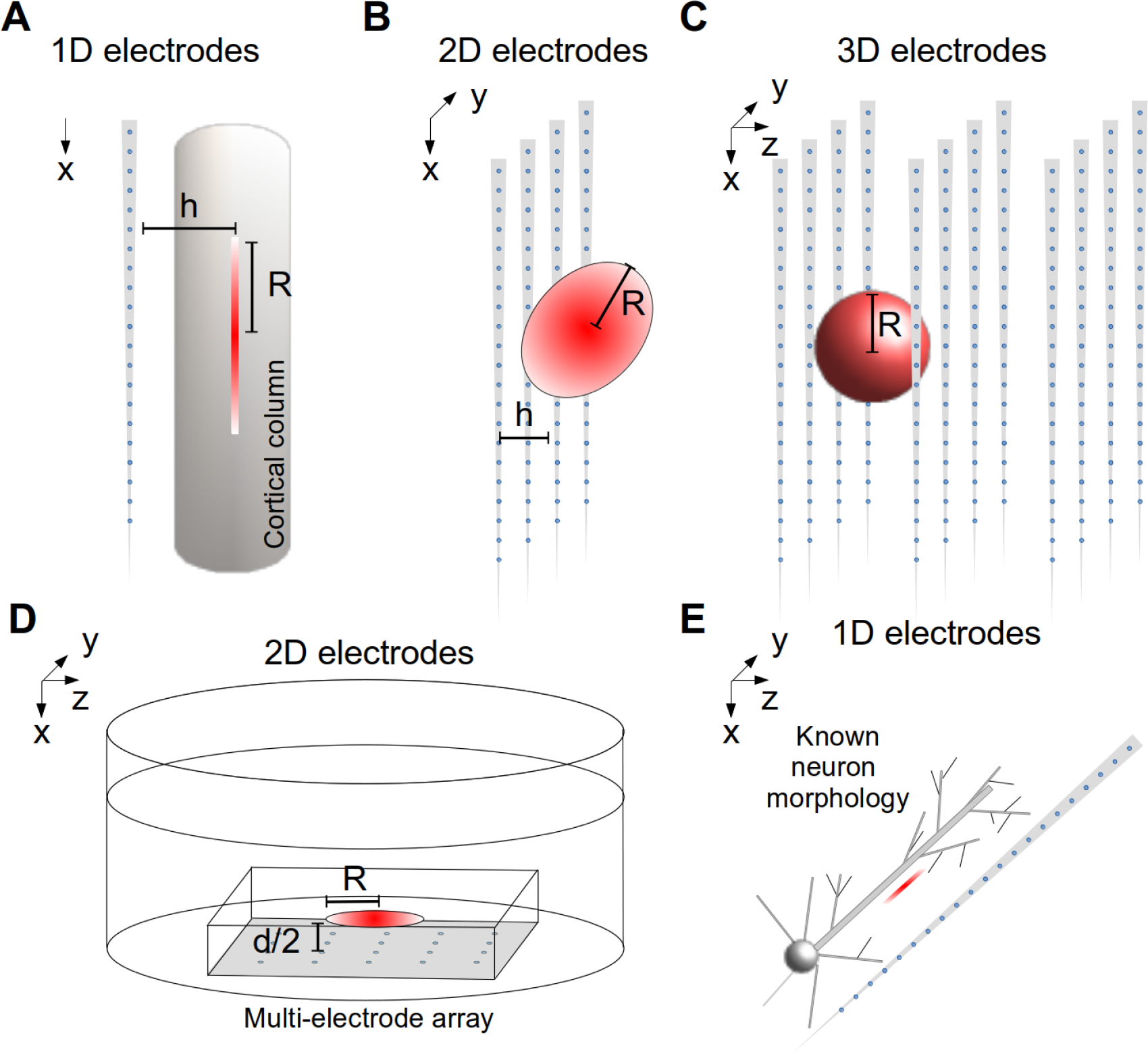
Overview of experimental contexts where `kCSD-python` is applicable. 1D setups such as A) laminar probes and equivalent; 2D setups, such as B) multishaft silicon probes, Neuropixel or SiNAPS probes, or D) planar MEA; 3D electrode setups, such as multiple multishaft silicon probes, Utah arrays, multiple electrodes placed independently in space with controlled positions, where the sources are assumed to come C) from tissue (kCSD) or E) from single cells with known morphology (skCSD). For description of parameters see Methods.

We also introduce some new diagnostic features which we found useful in CSD analysis and illustrate the application of the package and the new diagnostic tools in typical analysis workflow implemented as a Jupyter notebook (Kluyver et al., 2016) tutorial.

For reader convenience we first briefly restate the kCSD method (Potworowski et al., 2012; Chintaluri et al., 2021). Then in the Results section we introduce several new diagnostic tools included in the presented package. First, we added L-curve method for parameter selection. Then, to get informed on reconstruction accuracy we introduce *measurement uncertainty maps*, which show how measurement noise affects the estimated CSD, and *reliability maps*, which help build intuition on reliability of estimates for classes of possible sources for a given setup. We then briefly introduce the new package which is extensively illustrated in the provided tutorial in the Supplementary Materials. This jupyter notebook (Kluyver et al., 2016) tutorial, which is part of the toolbox, is provided to facilitate understanding and usage of the kCSD method. The tutorial enables the user to conduct analysis of the CSD using simulated surrogate data (electrodes positions and recordings) or actual recordings.

## 2 Methods

Kernel Current Source Density method uses kernel interpolation of measured potential to obtain V(x) in the whole space:

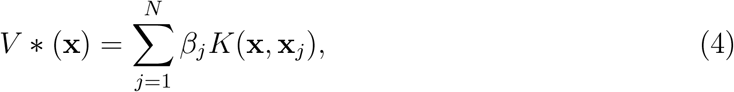

where **x**_*j*_, *j* = 1, …, *N*, are electrode positions, *K*(**x, x**^*′*^) is a symmetric kernel function. To avoid overfitting, correction for noise is made by minimizing prediction error

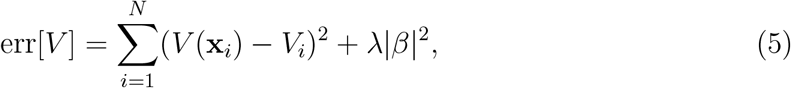

which gives

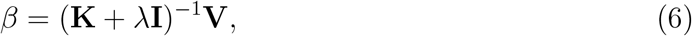

where **V** is the vector of measured potentials, *λ* is regularization parameter, and

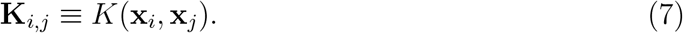

In the second step, we use function 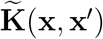, which we call cross-kernel, to estimate the CSD:

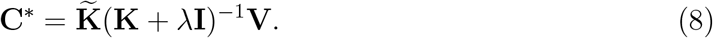

This procedure would work for arbitrary smoothing kernels **K** but in general it is difficult to identify the relevant corresponding 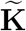, except in the simple cases. To circumvent this in kCSD we introduce a large basis of CSD sources spanning the region of interest, 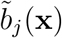, think of many little Gaussians, and corresponding basis sources in the potential space, *b*_*j*_(**x**), and construct both kernels from these basis sources (Potworowski et al., 2012; Chintaluri et al., 2021)

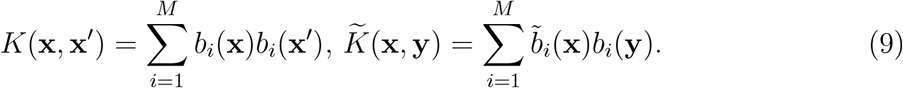

In these basis the CSD and the potential are given by

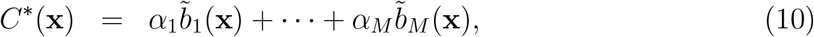

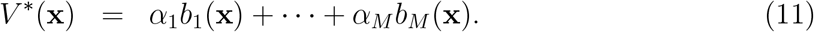

The challenges of the method are how to construct **K** and 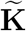, how to select the basis, the relevant parameters, and reliability of the estimation, some of which were introduced before (Potworowski et al., 2012; Chintaluri et al., 2021) and some we introduce here and illustrate with the provided package. For more details of kCSD method and the theory behind, see (Potworowski et al., 2012; Chintaluri et al., 2021).

## 3 Results

This paper introduces the `kCSD-python` package, an implementation of the kernel Current Source Density method (Potworowski et al., 2012) and its two variants ((Ness et al., 2015) and (Cserpan et al., 2017)). It is open source and available under the modified BSD License (3-Clause BSD) on GitHub (https://github.com/Neuroinflab/kCSD-python). It utilizes the continuous integration provided by Travis CI. It supports Python 3.9 version^1^ and has a bare minimal library requirements (`numpy`, `scipy` and `matplotlib`). It can be installed using the Python package installer (`pip`) or using Anaconda python package environment (`conda`). Details of the installation can be found in the package documentation at https://kcsd-python.readthedocs.io/en/latest/INSTALL.html.

The package contains a set of tools for kCSD analysis and to validate the results obtained from this analysis. To facilitate uptake of this resource, the package comes with extensive tutorials implemented in jupyter notebooks. These tutorials allow users to test different configurations of current sources and electrodes to see the method in action. The users can analyze their own data or explore the method with data generated *in silico*. These provisions illustrate the advantages and limitations of kCSD method to its users. The tutorials can also be accessed without any installation on a web browser via Binder (Project Jupyter et al., 2018). The package is extensively documented (https://kcsd-python.readthedocs.io) and includes all the necessary scripts to generate the figures in this manuscript.

An extensive tutorial overview of the `kCSD-python` package, is provided as an Appendix. Its goal is to show how to use `kCSD-python` to perform CSD analysis, how to apply the provided analysis tools, and to validate the results. We first consider a regular grid of ideal (noise-free) electrodes, where we compute the potentials from a known test source (the ground truth). We then use these potentials to reconstruct the sources which we compare with ground truth (Basic features). Then, we explore the effects of noise on the reconstruction and test the robustness of the method (Noisy electrodes). In the final part of the tutorial we look at how the errors in the estimation depend on the sources and the electrode configuration by testing the effects of broken electrodes on reconstruction (Broken electrodes).

Supplementary Fig. 11 shows error propagation maps, which we introduce below, for 1D regular grid of 12 electrodes. Supplementary Fig. 12 shows an example of 3D kCSD source reconstruction. Supplementary Fig. 13 shows an example of skCSD reconstruction (single cell kernel CSD) which corresponds to Fig. 8 from Cserpan et al., 2017. The simulation, reconstruction and visualization have all been re-implemented in Python.

We also provide some pre-computed examples of CSD estimations using our library for users to explore. We provide these as `.pdf` files available at http://bit.ly/kCSD-suppleme Here, the files `small_srcs_3D_all.pdf` and `large_srcs_3D_all.pdf` show 100 example setups of 3D kCSD reconstructions from small and large sources. Similarly, files `small_srcs_all.pdf` and `large_srcs_all.pdf` show 100 example setups of 2D kCSD reconstructions from small and large sources. “Smallness” and “largeness” of sources is defined by the ratios of typical spatial scales of the source with respect to interelectrode distances.

### 3.1 Parameter selection

An important part of kCSD estimation is selection of parameters, in particular the regularization parameter, *λ*, but also the width of the basis source, *R*. Previously we proposed to use cross-validation (Potworowski et al., 2012). Here we introduce L-curve approach (P. Hansen, 2010; Kropf and Shmuel, 2016) for regularization. Both these methods are implemented in `kCSD-Python`.

#### Cross-validation

To select parameters using cross-validation (Potworowski et al., 2012) we consider a range of parameter values, *λ∈* [*λ*_0_, *λ*_1_]. For any test value *λ* we select an electrode *i* = 1, …, *N* and ignore it. With eq. (4) we build a model from remaining measurements, 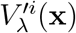, and use it to predict the value at the ignored electrode, 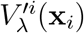.

Here

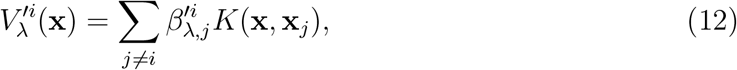

where the minimizing vector

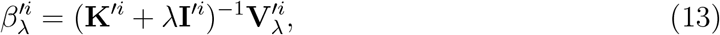

and where ^*′i*^ means *i*-th column and row are removed from the given matrix (or vector). We repeat this for all the electrodes *i* = 1, …, *N* and compare predictions from the remaining electrodes against actual measurements:

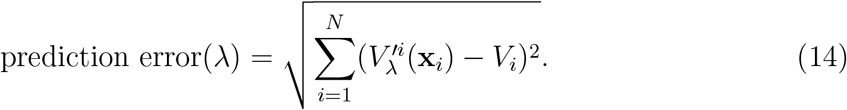

For final analysis, *λ* giving minimum prediction error is selected. It is worth checking if the global minimum is also a local minimum. If the *λ* selected is one of the limiting values this may indicate that extending the range of *λ* might result is more optimal result or that the problem is ill-conditioned, for example too noisy, and we are either underfitting or overfitting, as we discuss below for the L-curve. As a rule of thumb, the range of tested *λ* parameters should cover eigenvalues of the (7).

#### L-curve

Consider the error function, Eq. (5), which we minimize to get the regularized solution, *V*_*λ*_ = **K***β*_*λ*_ It is a sum of two terms we are simultaneously minimizing, prediction error

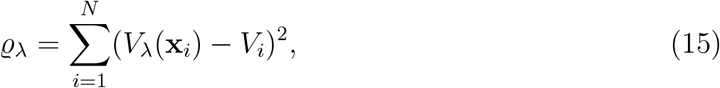

and the norm of the model

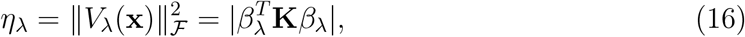

weighted with *λ*. Taking *λ* = 0 is equivalent to assuming noise-free data. In this case we are fitting the model to the data, in practice, overfitting. On the other hand, taking large *λ* means assuming very noisy data, in practice ignoring measurements, which results in a flat underfitted solution. Between these extremes there is usually a solution such that if we decrease *λ*, the prediction error, *ϱ*, slightly decreases, while the norm of the model, *η*, increases fast, and if we increase *λ*, the prediction error, *ϱ*, increases fast, while the norm of the model, *η*, slightly decreases; see Fig. 2D.

**Figure 2.**
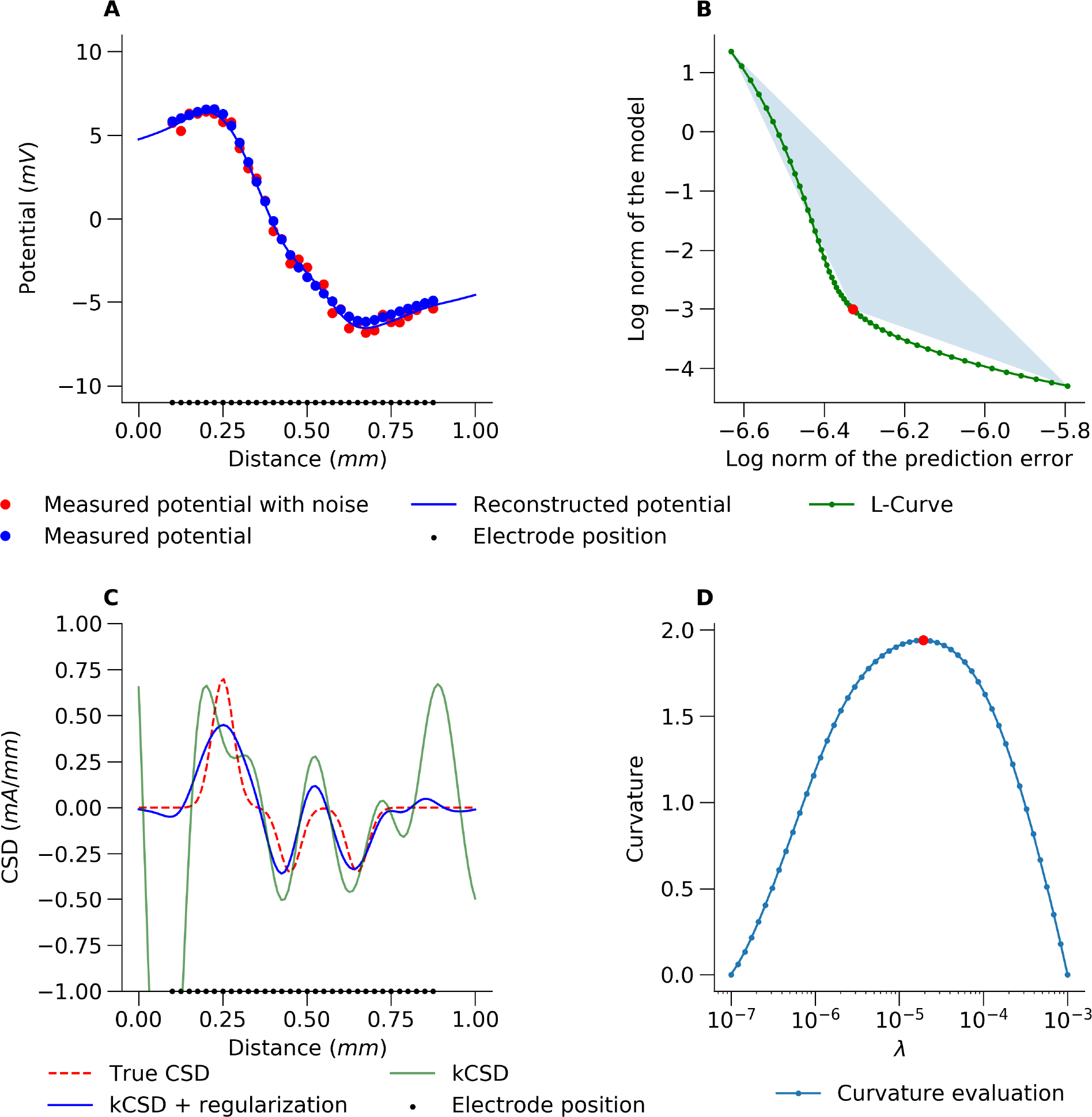
An example of the L-curve method for estimating kCSD parameters. A) The red points represent the potential used for CSD reconstruction. The black points show the electrode positions. The ground truth is shown in panel C with the red dashed curve. The measurement was simulated by adding small random noise to all the electrodes (32 values taken from a uniform distribution). The blue line shows a kernel interpolation of the potential which is the first step of kCSD method. B) L-curve plot for a single R parameter. The apex of the L-curve is numerically computed from the oriented area of directed triangles connecting the point on the L-curve with its two ends. C) Comparison of the true CSD and kCSD reconstruction for parameters obtained with L-curve regularization. D) Estimation of L-curve curvature with triangle method (see the Methods).

This is apparent when the prediction error and the norm of the model are plotted in the log-log scale. This curve follows the shape of the letter L, hence the name L-curve (P. C. Hansen, 1992). Several methods have been proposed to measure the curvature of the L-curve and to identify optimal parameters (P. C. Hansen, Jensen, and Rodriguez, 2007). In `kCSD-python`, we have implemented the triangle area method proposed by Castellanos, Gómez, and Guerra, 2002. To distinguish between convex and concave plot, clockwise directed triangle area is measured as negative Fig. 2.D shows this estimated curvature for our example as a function of *λ*.

To illustrate this method in the context of CSD reconstructions, we study an example of 1D dipolar current source with a split negative pole (sink; see Fig. 2.C, True CSD, red dashed line). We compute the potential at 32 electrodes (Fig. 2.A, C, black dots) with additive noise at every electrode. Notice that if we want to interpret the recorded potential directly (Fig. 2.A, red dots) it is difficult to discern the split sink. Fig. 2.D shows the estimated curvature for our example as a function of *λ*. The optimal value of *λ* is found by maximizing the curvature of the log-log plot of *η* versus *ϱ*, Fig. 2.B. The red dot in Fig. 2.B, D, indicates the ideal *λ* parameter for this setup obtained through the L-curve method.

### 3.2 Reconstruction accuracy

With kCSD procedure one can easily estimate optimal CSD consistent with the obtained data. However, so far we have not discussed estimation of errors on the reconstruction. Since the errors may be due to a number of factors — the procedure itself, measurement noise, incorrect assumptions — one may approach this challenge in different ways.

First, to understand the effects of the selected basis sources and setup, one may consider the estimation operator 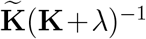 and the space of solutions it spans. This space is given by the eigensources, introduced and described thoroughly in (Chintaluri et al., 2021). The orthogonal complement of this space in the original estimation space, spanned by 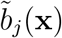 basis functions, is not accessible to the kCSD method. The study of eigensources facilitates understanding which CSD features can be reconstructed and which are inaccessible.

Second, to consider the impact of the measurement noise on the reconstruction, for any specific recording consider the following model-based procedure. Reconstruct CSD from data with optimal parameters. Compute potential from estimated CSD. Add random noise to each computed potential. The noise could be estimated from data, either as a measure of fluctuations on a given electrode for a running signal, or from variability of evoked potentials. Then, for any realization of noise, compute estimation of CSD. The pool of estimated CSD gives estimation of the error at any given point where estimation is made.

This computation can be much simplified by taking advantage of the linearity of the resolvent, 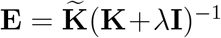. Then, the *i*-th column (**E**_*i*_) represents contribution of unitary change of *i*-th measured potential (the *i*-th element of the vector **V**) to the estimated CSD (**C**^*\**^). As the contribution is proportional to the change, the column can be considered an *Error Propagation Map* for *i*-th measurement (Fig. 3.A). Note that these vectors (the columns of resolvent, **E**_*i*_) also happen to form another basis of the solution space, an alternative to the basis of eigensources.

**Figure 3.**
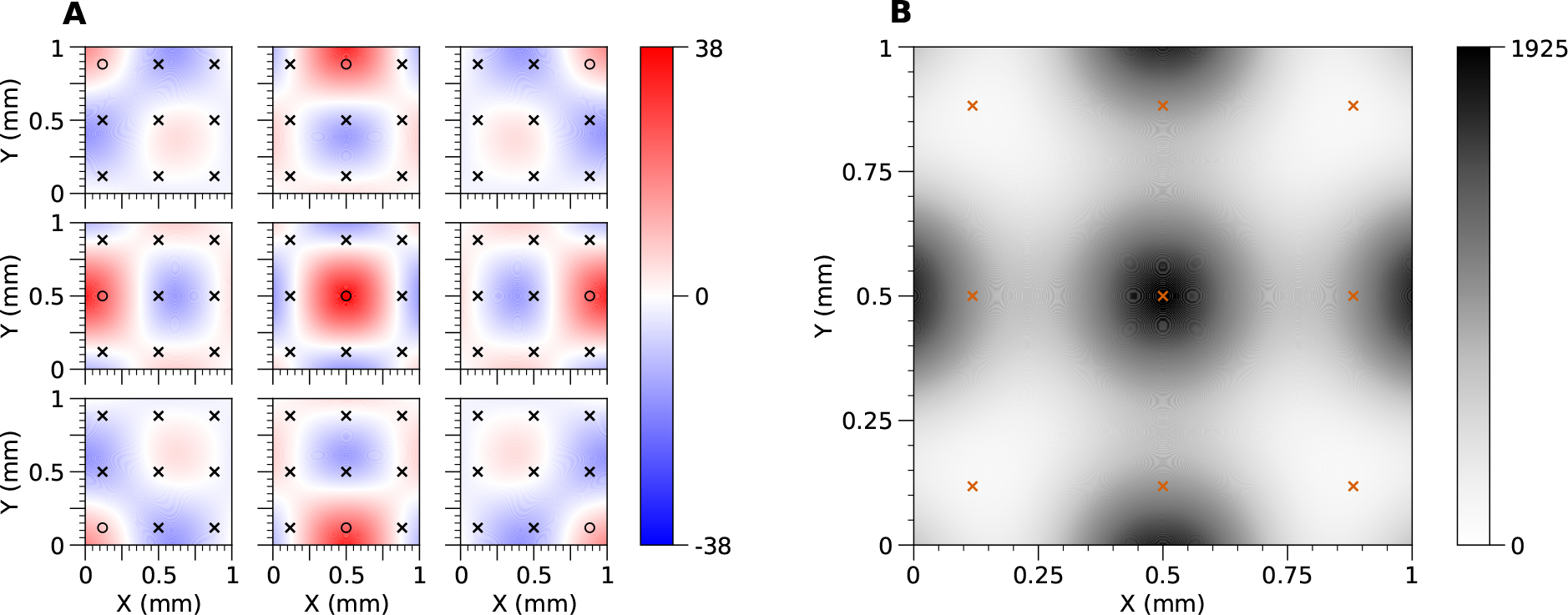
A) Error propagation maps for 3*×*3 regular grid of electrodes. Every panel represents the contribution of the potential measured at the corresponding electrode marked with a black circle (○) to the reconstructed CSD. Every other electrode is marked with a black cross (*×*). B) Map of CSD measurement uncertainty for 3*×*3 regular grid of electrodes. The CSD measurement uncertainty is represented by variance of the CSD reconstruction caused by the uncertainty in measurement of the potentials. It is assumed that measurement errors for electrodes are mutually independent and follow standard normal distribution (*ε*_*i*_ *∼ N* (0, 1)). Location of electrodes is marked with red crosses (*×*). Fig. 4 shows example reliability map for the case of 10×10 electrode distribution.

If *ε*_*i*_ is an error of *i*-th measurement, then its contribution to **C**^*\**^ is *ε*_*i*_**E**_*i*_. Moreover, if the measurement errors follow multivariate normal

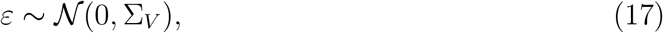

Then

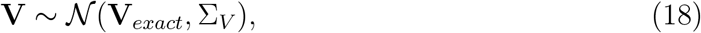

and the estimated CSD also follows multivariate normal

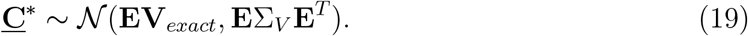

The diagonal of **E**Σ_*V*_ **E**^*T*^ represents a *map of CSD measurement uncertainty* (uncertainty attributed to the noise in the measurement, Fig. 3.B)^2^.

Third, one can study reconstruction accuracy for a meaningful family of test functions. This could be the Fourier modes for rectangular regions or a collection of Gaussian test functions, centered in different places, of single or multiple radii. For each of these test functions one would compute the potential, perform reconstruction, and compare the results with the original at every point. Finally, one could average this information over multiple different test sources computing a single *Reliability Map*, which we now introduce.

#### Reliability maps

Assume the standard kCSD setup, that is a region *ℛ⊂*ℝ^*n*^ where we want to estimate the sources, set of electrode positions, **x**_*i*_, and perhaps additional information, such as morphology for skCSD Cserpan et al., 2017. We now want to characterize predictive power of the combination of our setup and our selected basis, 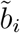. To do this we select a family of test functions, *C*^*i*^(**x**), for example Gaussian test functions, centered in different places, of multiple radii, or products of Fourier modes, etc. Then, for each *C*^*i*^ we compute *V* ^*i*^=*𝒜C*^*i*^ by forward modeling, generating a surrogate dataset. Next, we apply the standard kCSD reconstruction procedure obtaining estimation of the tested ground truth, 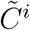. We can then compute reconstruction error using point-wise modification of Relative Difference Measure (RDM) proposed by (Meijs et al., 1988):

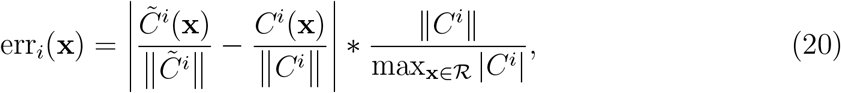

where *i* = 1, 2, … enumerates different ground truth profiles. A simple measure of reconstruction accuracy is then given by the average over these profiles:

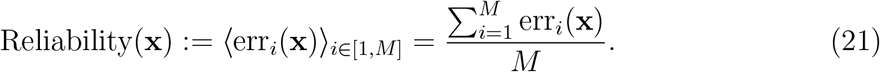

The class of functions used were the families of small and large sources mentioned above. We used eight mirror symmetries of the grid in computation.

We can use reliability map as another source of information about the precision of reconstruction, which is shown in Fig. 5. In A) we show some dipolar source which is used to compute the potential on a grid of electrodes shown in B). Fig. 5.C) shows reconstructed sources superimposed on reliability map. Panel D) shows the difference between the ground truth and reconstruction. Note that plots such as shown in panel A) and D) are feasible only for simulated or model data, where we know actual sources and use them to validate the method. On the other hand, plots shown in panel B and C represent what can be routinely computed for experimental data. This functionality is implemented in `kCSD-python` and illustrated in the provided tutorial.

**Figure 4.**
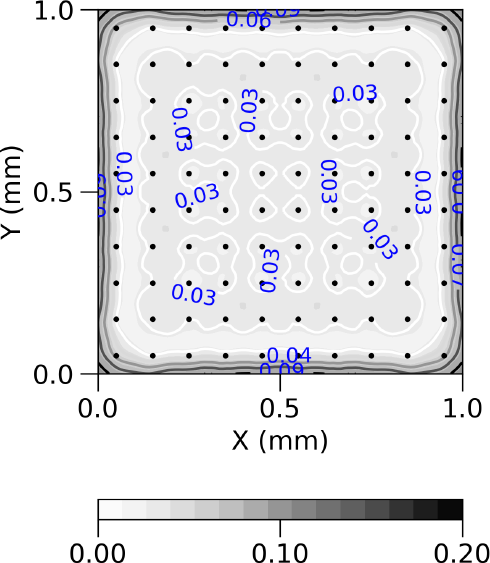
Reliability map created according to formula (20) and (21) for 10×10 regular grid of electrodes with noise-free symmetrized data. Black dots represent locations of contacts used in the study. Values on the map can be interpreted as follows: the closer to 0, the higher reconstruction accuracy might be achieved for a given measurement condition.

**Figure 5.**
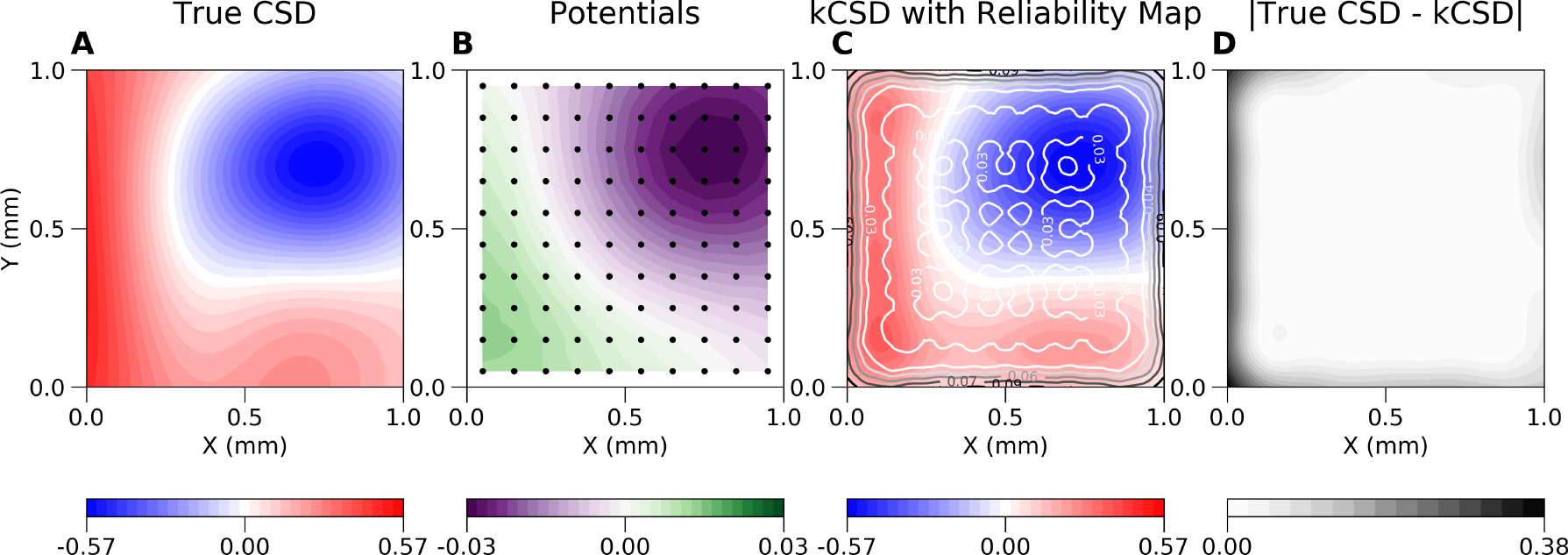
Example use of reliability maps. A) Example dipolar source (ground truth) which is used to compute the potential on a grid of electrodes shown in B). C) shows reconstructed sources superimposed on reliability map. D) shows the difference between the ground truth and the reconstruction.

Another interesting question is the effect of broken or missing electrodes on the reconstruction. Formally one can attempt kCSD reconstruction from a single signal but it is naive to expect much insight this way. It is thus natural to ask what information can be obtained from a given setup and what we lose when part of it becomes inaccessible.

Fig. 6 shows the effect of removing electrodes on the reconstruction. Fig. 6.A shows average error of kCSD method across many random ground truth sources for a regular grid of 10×10 electrodes. Fig. 6.B to D show the increase of average reconstruction error as we remove 5 (B), 10 (C) and 20 (D) contacts. To emphasize the errors we show the difference between the reliability map for the broken grid minus the original one. Note the different scales in plots B–D versus A The consecutive rows show similar results when only small sources were used (E–H), or only large sources were used (I–L). Random sources in Fig. 6.A are both small and large sources (mentioned in the Results). This shows, among others, as we explained, that the reliability maps depend on the test function space, however, we feel they are more intuitive to understand than the individual eigensources spanning the solution space (Chintaluri et al., 2021).

**Figure 6.**
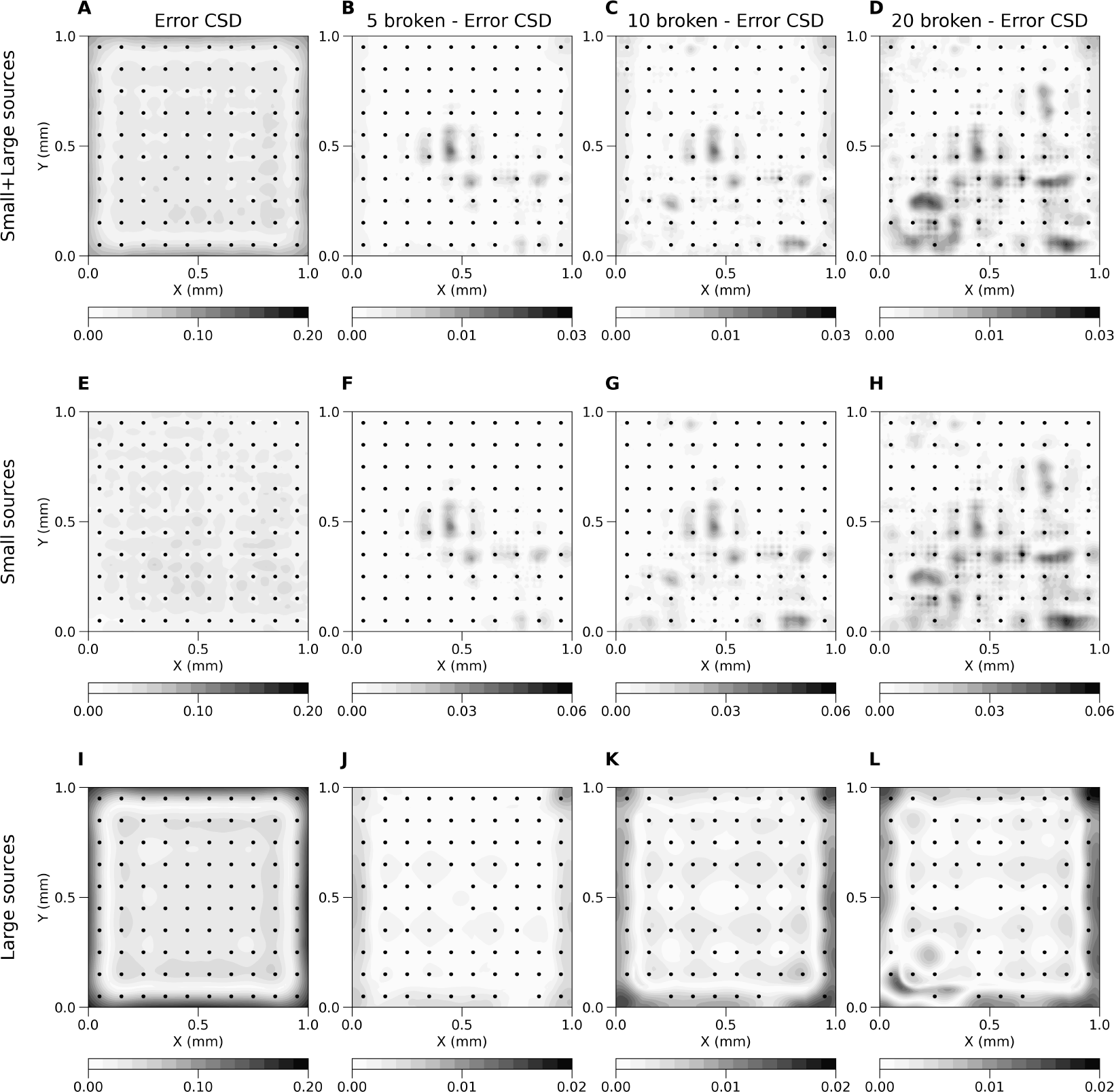
Average error (eq. 20) of kCSD method across random small and large (A), only small (E) and only large (I) sources for regular 10×10 electrodes grid and the same grid with broken 5 (B, F, J), 10 (C, G, K) and 20 (D, H, L) contacts. Plots (B, C, D, F, G, H, J, K, L) show difference between average error for regular grid and grid with broken contacts. Estimation was made in noise free scenario, *R* parameter selected in cross-validation. Black dots represent locations of contacts used in the study.

## 4 Discussion

In the present work we introduced a new Python package for kCSD estimation, `kCSD-python`. This toolbox implements the basic functionality of source estimation with regularization. We introduced here L-curve approach as an approach to parameter selection alternative to cross-validation. This might be particularly important if regularization is attempted for high-density setups, such as multi-electrode arrays with thousands of channels. We also proposed several ways of diagnosing reconstruction errors, specifically Error Propagation Maps and Reliability Maps, which can build intuition regarding veracity of reconstructions obtained in given experimental contexts. Finally, we provided a brief tutorial to the new package (see Supplementary Material) illustrating its main functions. All the figures in this paper showing CSD and LFP were computed with this package and source files are provided. In this section we discuss several issues related to CSD analysis in general and kCSD in particular.

### The benefits of CSD estimation over direct LFP analysis

Extracellular potentials provide valuable insight about the functioning of the brain. Thanks to recent advances in multielectrode design and growing availability of sophisticated recording probes we can monitor the electric fields in the nervous system at thousands of sites, often simultane-ously. One may wonder if this increased resolution makes CSD analysis unnecessary. In our view, as we have discussed many times, it is not so. The long range nature of electric potential means that even a single source contributes to every recording. Thus in principle we should always expect strong correlation between neighboring sites. However, if the separation between the electrodes becomes substantial, on the order of millimeters, the level of correlation between the recordings on different electrodes will decrease. This is because each electrode effectively picks up signals from a different composition of sources. Even if some are shared they are overshadowed by others which may lead to small interelectrode correlations. Still, our experience shows that significant correlations can be observed in the range of several millimeters (Łęski, Wójcik, et al., 2007; Katzner et al., 2009; Hunt et al., 2011; Kajikawa and Schroeder, 2011) which is consistent with computational models (Lindén, Tetzlaff, et al., 2011; Łęski, Lindén, et al., 2013). Fundamentally, the LFP profile is different from CSD profile, and may significantly distort or hide features of importance. For example, Fig. 2.C shows the source composed of three gaussians, while direct inspection of LFP suggests a simple dipole.

In view of these facts we argue that it is always beneficial to attempt kCSD analysis. The caveat is not to believe the reconstructed CSD blindly but always interpret it against known anatomical and physiological knowledge supported by the tools such as provided in the present work (eigensources, reliability maps, etc). This could be combined with other decomposition methods if desired which may give more physiologically interpretable results, in particular we have had very good results using independent component analysis, which we recommend (Łęski, Kublik, et al., 2010; Głąbska et al., 2014).

### Approaches to CSD estimation

Several procedures for CSD estimation were introduced over the years and are still in use. The first approach, which probably still dominates today, was introduced by Walter Pitts in 1952 (Pitts, 1952) and gained popularity after Nicholson and Freeman adopted it for laminar recordings (Nicholson and Freeman, 1975). This was a direct numerical approximation to computation of the second derivative in the Poisson equation (1). Only minor improvements were introduced over the years to stabilize estimation (Rappelsberger, Pockberger, and Petsche, 1981) or handle boundaries (Vaknin, DiScenna, and Teyler, 1988). The first major conceptual change was introduced by Pettersen et al., 2006 who introduced model-based estimation of the sources. Their idea was to assume a parametric model of sources, for example, spline interpolated CSD between electrode positions, and using forward modeling to connect measured potentials to model parameters. This model-based approach was generalized by Potworowski et al., 2012 who proposed a non-parametric kernel Current Source Density method which is implemented in the presented toolbox.

Apart from these main approaches several CSD variant methods were proposed. For example, one may interpolate the potential first before applying traditional CSD approach, or the opposite, interpolate traditionally estimated CSD. Although in some cases the obtained results may look close to those obtained with kCSD, we do prefer kernel CSD approach due to the underlying theory which facilitates computation of estimation errors but also yields a unified framework for handling underlying assumptions, noisy data and irregular electrode distributions. In our view approaches combining ad hoc interpolation with numerical derivatives conceptually and computationally are less convincing to iCSD and kCSD and we would not recommend them.

### Models of tissue

Throughout this work and in the toolbox we assumed purely ohmic character of the tissue. This has been debated in recent years (Bédard and Destexhe, 2011; Riera et al., 2012; Gratiy et al., 2017) and it is true that more complex biophysical models of the tissue, taking into account frequency dependent conductivity or diffusion currents, would influence the practice of source reconstruction or its interpretation. However, the available data indicate that in the range of frequencies of physiological interest these effects are small. While one should keep eyes open on the new data as they become available and keep in mind the different possible sources which may affect the reconstruction or interpretation, we believe that the traditional view of ohmic tissue is an adequate basis for typical experimental situations and going beyond that would probably require additional dedicated measurement for the experiment at hand which may not always be feasible. For example, as we discussed in (Ness et al., 2015), the specimen variability of the cortical conductivity in the rat is much bigger than the variability between different directions within a given rat (Goto et al., 2010). This means that unless we have conductivity measurements for our specific rat we are probably making smaller error assuming isotropic conductivity than taking different values from literature. We feel there is not enough data to justify inclusion of more complex terms in the standard CSD analysis to be applied throughout the brains and species.

In this manuscript and in the `kCSD-python` toolbox we also assumed constant conductivity (with exception of MoI case below). We are convinced this is a reasonable approximation for typical depth recordings. In general, however, this approximation needs to be justified or alternative models of tissue need to be considered. In principle, the kCSD method can be applied for a variety of tissue models as long as the basis potentials can be computed from the basis sources while incorporating the geometric and conductivity changes.

For example, Ness et al., 2015 considered a cortical slice placed on a microelectrode array (MEA) in which they included the geometry of the slice and modeled saline-slice interface with changing conductivity in the forward model. They found that Method of Images (MoI) gives a good approximation to the full solution obtained using finite-element model (FEM). This approximation was incorporated within the kCSD method as MoIkCSD variant and is available in the `kCSD-python` package.

It is possible to generalize kCSD to reconstruct sources from recordings of multiple electrical modalities — LFP, ECoG, EEG. In this case one needs to include the head geometry and the changing tissue properties within the forward model and in the kCSD method. The anisotropic (white matter tracts) and inhomogeneous (varying between skull, cerebro-spinal fluid, gray matter and white matter) electrical conductivity changes can be approximated using data obtained with imaging techniques such as MRI, CT or DTI. Such sophisticated head models require numerical solutions such as finite element modeling (FEM) to compute the basis potentials from the basis sources. We are currently working on this approach to make it generic for any animal head and to eventually utilize it as a source localization method for human data, for example, to localize foci of pharmacologically intractable epilepsy seizures in humans. We call this extension kernel Electrical Source Imaging (kESI).

### High density microelectrode recordings

One of the trends clearly observed in modern neurotechnology is the drive towards increasing the number of sensors and their density (Buzsáki, 2004; Berdondini et al., 2005; Frey et al., 2009; Hottowy et al., 2012; Jun et al., 2017; Angotzi et al., 2019), for *in vitro* and *in vivo* studies. While it seems that a better resolution for recording spiking activity of multiple cells is the main goal, also more precise stimulation and field potentials monitoring are targeted (Hottowy et al., 2012; Ferrea et al., 2012; Bakkum et al., 2013). Such massive high density data from thousands of electrodes should greatly increase insight into the studied systems and significantly improve results of CSD reconstructions. There are two obstacles to fully benefit from kCSD analysis of data from these new systems. First, kCSD involves inversion of the kernel matrix which is quadratic in the number of electrodes. Combined with cross-validation the necessary matrix operations quickly become overwhelming. This can be mitigated in a number of ways, by subsampling the data, approximate inversions, and by switching from cross-validation to L-curve method, but the challenge remains. This is the easy problem. The difficult problem is physical. As we move away from a source its contribution to the recorded potential goes down. In consequence, since the present version of kCSD uses all recordings to estimate every source, when using remote signals to estimate local source, we obtain mainly contributions from noise. In effect we get a very reliable estimation of sources varying slowly in space but the sources changing fast in space are treated as noise and silenced by the regularization. To take full advantage of these data a different approach may give better results. One possibility could be a multiscale approach where one would perform reconstructions in small windows in multiple scales to optimally reconstruct multiscale features of the source distribution. The challenge would be to efficiently and correctly stitch them together.

### Parameter selection

In the Results section we discussed our strategy for data-based parameter selection using cross-validation or L-curve. Often, we need to tune not just *λ* but also other parameters. For example, for Gaussian basis sources we may want to decide on the width of the Gaussian used, *R*. To obtain the optimal set of parameters in that case we compute the curvature of the L-curve or the cross-validation error for some ranges of parameters considered and select parameters corresponding to the maximum curvature / minimum error in the parameter space. This is a simplification of the proposition by Belge, Kilmer, and Miller, 2002 which in practice we found very effective.

As an example, in Fig. 7 we show results of such a scan for the problem shown in Fig. 2. The range of *λ* to be considered can be set by hand but by default we base it on the eigenvalues of *K*. The smallest *λ* is set as the minimum eigenvalue of *K* which here was around 1e-10. We set maximum *λ* at standard deviation of the eigenvalues, which here was around 1e-3. The range of *R* values studied was from the minimum interelectrode distance to half the maximum interelectrode distance. Note that for very inhomogeneous distributions of electrodes this approach may be inadequate.

**Figure 7.**
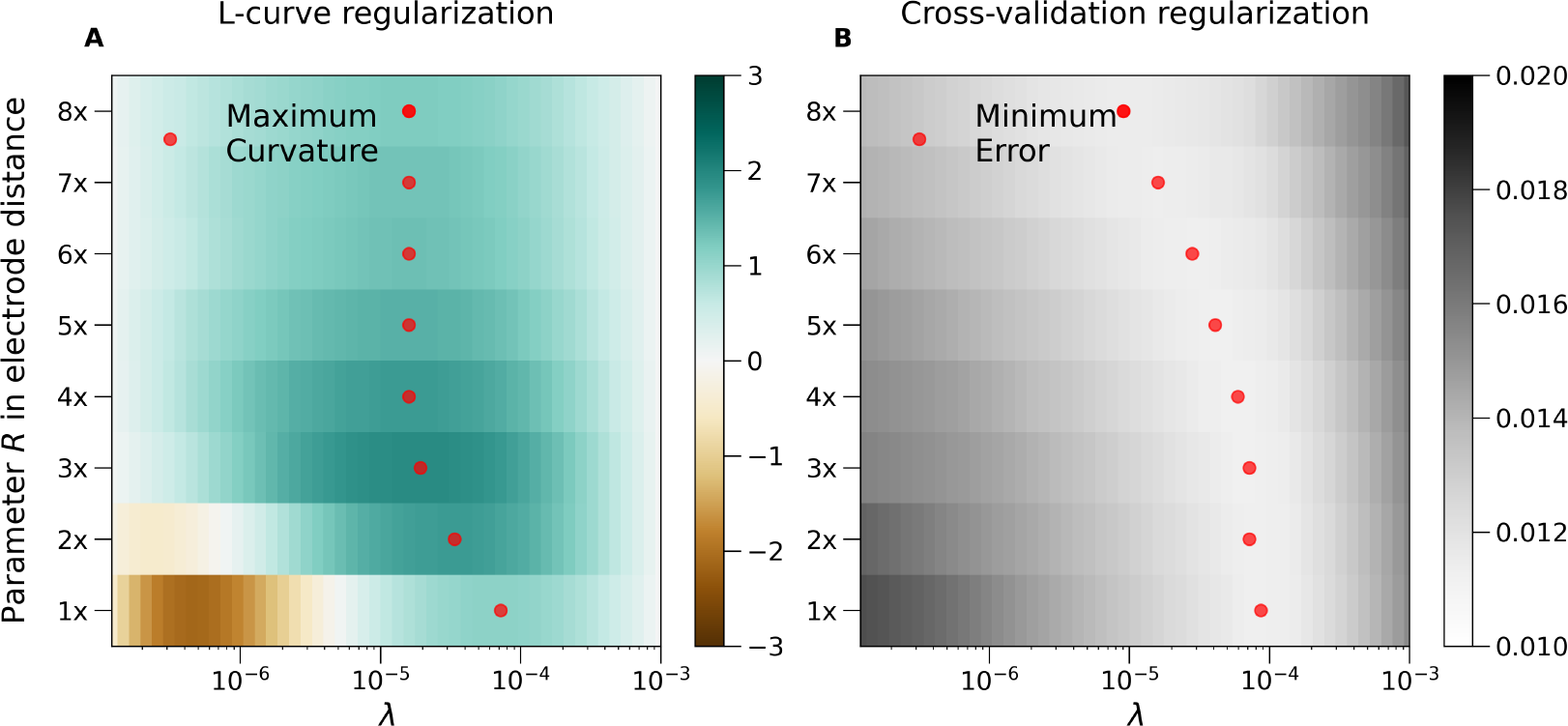
L-curve curvature (left) and CV-error (right) for the problem studied in Fig. 2. Observe that in both cases there are ranges of promising candidate parameter pairs, *R, λ*, which can give good reconstruction given the measured data. Red dots shows local extrema for each value of *R* fixed. See text for discussion of this effect.

**Figure 8.**
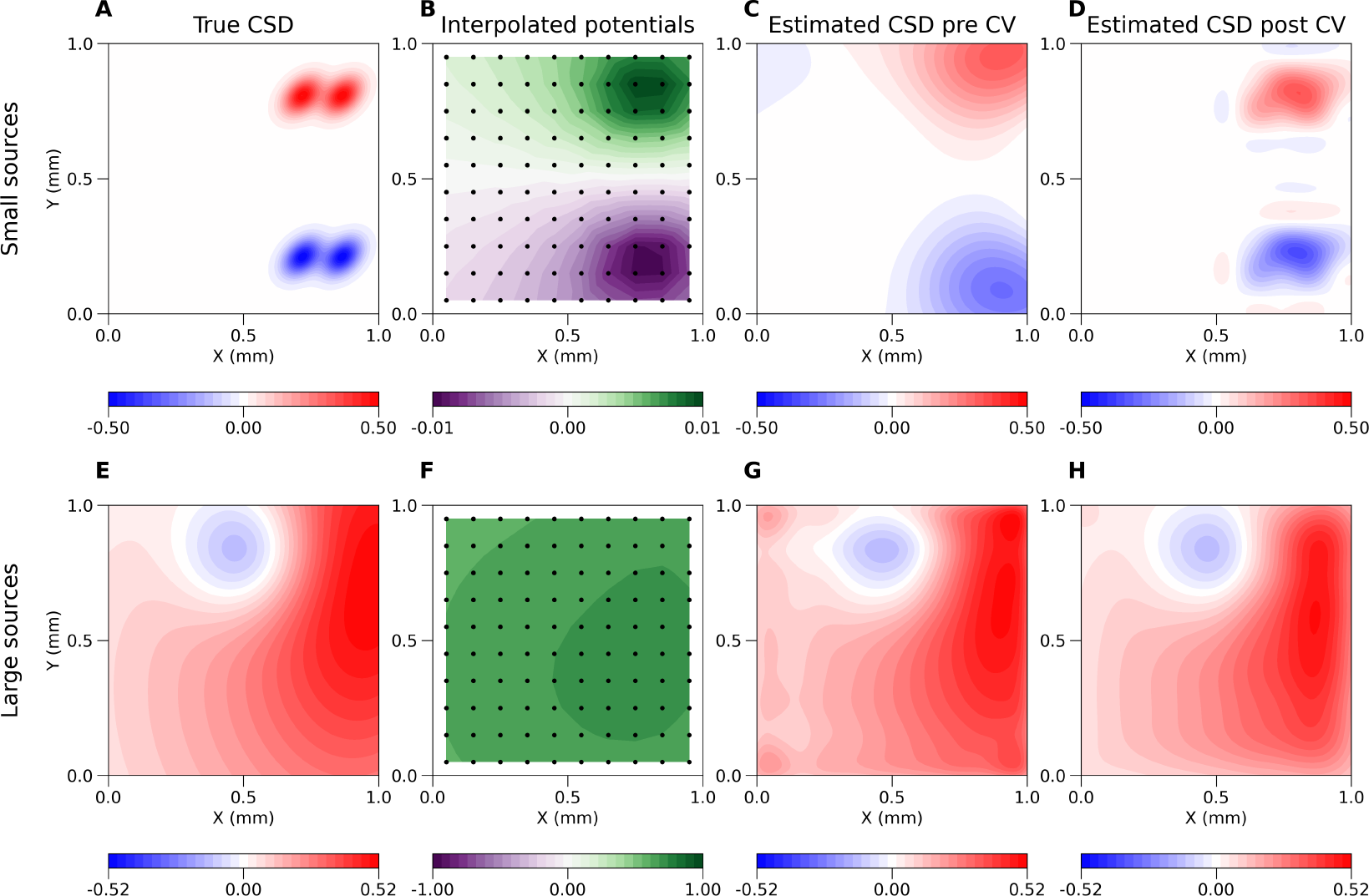
Basic features tutorial A) Shows the ground truth (True CSD), here, two-dimensional small Gaussian current sources, for the CSD seed of 15. B) The interpolated potentials generated by this current source are shown, the electrodes are displayed as black dots. C) CSD estimated with kCSD using the potentials from the electrode positions, without cross-validation. D) Same as C but cross-validation was used. E-H) Analogous to A–D, except large Gaussian current sources for seed 6 were used.

What we find is that apart from a global minimum in *R, λ* space there is a range of *R* values fixing which we can find optimal *λ*(*R*) which leads to very close curvatures / CV-errors / estimation results. What happens is that within some limits we may achieve similar smoothing effects changing either *λ* or *R*. Bigger *λ* means more smoothing, but bigger *R* means broader basis functions and effectively also smoother reconstruction space. This is why the CV-error and curvature landscapes are relatively flat, or have these marked valleys observed in Fig. 7. This effect supports robustness of the kCSD approach.

### The sources of error in kCSD estimation and how to deal with them

Kernel CSD method assumes a set of electrode positions and a corresponding set of recordings. Additionally, single cell kCSD requires morphology of the cell which contributed to the recordings and its position relative to the electrodes. Each of these may be subject to errors.

In analysis, we assume that the electrode positions are known precisely. This is a justified assumption in case of multishaft silicon probes or integrated CMOS-MEA but not necessarily when multiple laminar probes are placed independently within the brain or for many other scenarios. For example, for SEEG electrodes in human patients used in presurgical evaluation we expect the localization errors due to the workflows used clinically to be significant. We do not provide dedicated tools to study the effects of misplaced electrodes on the reconstructed CSD, however, this can be achieved easily with the provided package if needed. The location of the cell with respect to the electrodes is much more questionable, especially in 3D. Nevertheless, the necessary data to perform skCSD today are too scarce to start addressing these issues.

On the other hand we do assume that the recordings are noisy and we use regularization to counteract the effects of noise. We have no mechanism to differentiate between electrodes with varying degrees of noise to compensate this differently. However, we observed that for cases with very bad electrodes, similar results are obtained for analysis of complete data and for analysis of partial data with bad electrodes removed from analysis. The difference was in *λ* selected which was larger when broken electrodes were included in the analysis. Depending on situation, if there is a big difference in the noise visible in different channels, an optimal strategy may be to discard the noisy data and perform reconstruction from the good channels only, which kCSD permits. In the end, data analysis remains an art and a healthy dose of common sense is always recommended.

The main limitation of the method itself lies in the character of any inverse problem. Here it means that there is an infinite number of possible CSD distributions each consistent with the recorded potential. It is thus necessary to impose conditions which allow unique reconstruction and this is what every variant of CSD method is about. In kCSD this condition is minimization of regularized prediction error. In practical terms one may think of the function space in which we are making the reconstruction. This space is span by the eigensources we discussed before (Chintaluri et al., 2021). We feel it is useful to consider both this space as well as its complement, that is the set of CSD functions whose contribution to every potential is zero. This can facilitate understanding of which features of the underlying sources can be recovered and which are inaccessible to the given setup. While for the most common regular setups, such as rectangular or hexagonal MEA grids or multishaft probes, intuitions from Fourier analysis largely carry over, in less regular cases this quickly becomes non-obvious.

To facilitate intuition building in the provided toolbox we include tools to compute the eigensources for a given setup. We also proposed here reliability maps, heuristic tools to build intuition regarding which parts of the reconstructed CSD can be trusted and which seem doubtful. These reliability maps are built around specific test ground truth distributions and some default parameters facilitating validation for any given setup are provided. Due to the open source nature of the provided toolbox more complex analysis is possible if the setup or experimental context require that.

## Acknowledgements

The Python implementation of kCSD was started by Grzegorz Parka during Google Summer of Code project through the International Neuroinformatics Coordinating Facility. Jan Mąka implemented the first Python version of skCSD class. The study received funding from the Polish National Science Centre’s grants (2013/08/W/NZ4/00691) and (2015/17/B/ST7/04123). The authors declare no conflict of interest.

## Appendix

### kCSD-python package tutorial

In this section we first illustrate the use of kCSD package for CSD reconstruction in the simplest case of a regular 2D square grid. This is a simplified version of a slice on a microelectrode array (Ness et al., 2015), or a planar silicone probe within the brain, where we assume constant conductivity in the whole space. In the following sections we show how we validate our methods and what kind of diagnostics we find useful in the analysis of experimental data. This tutorial is available as a jupyter notebook and can also be accessed through a web-browser without installation. For more details, see ‘Overview of kCSD-python package’ in the Discussion.

### Basic features

We start with the basic CSD estimation on a regular grid. First, we define a region of interest. Then, using predefined test functions for the current sources, we place a ground truth current source in this region. We define the distribution of electrodes. Assuming ideal electrodes, we compute the potential generated by the selected current sources as measured at the electrodes. Given these potentials and the electrode locations we estimate the current source density using kCSD. As a final step, we perform cross-validation to avoid overfitting. Since we know the ground truth used to generate the potentials that were used in the kCSD estimation, we can compare the ground truth to the estimate and see the reconstruction accuracy.

### Defining region of interest

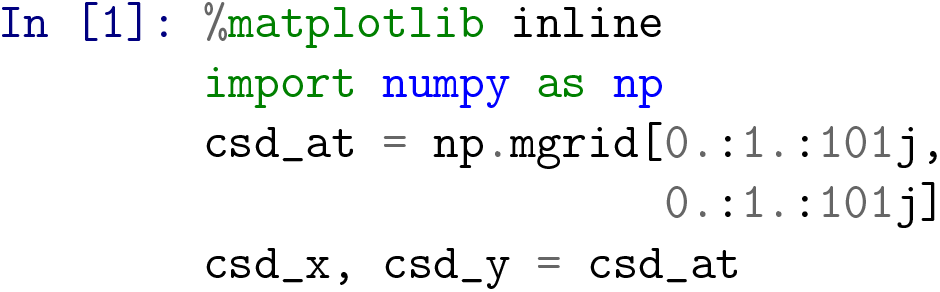

We define the region of interest between 0 and 1 in the `xy` plane with a resolution of `101` points in each dimension. We will assume the distance is given in *mm*, so we want to perform a reconstruction on a square patch of 1*mm*^2^ size.

#### Setting up the ground truth

The `kCSD-python` library provides functions to generate test sources which can be imported from the `csd_profile` module. Here we use the `gauss_2d_small` function to generate two-dimensional Gaussian sources which are small in the scale set by the interelectrode distance. The other implemented option for two-dimensional test sources is the `gauss_2d_large` function. To generate the exact same sources in each run we must invoke this function using the same random seed which is stored in the `seed` variable. For simplicity, these current sources are static and do not change with time. We visualize the current sources as a heatmap.

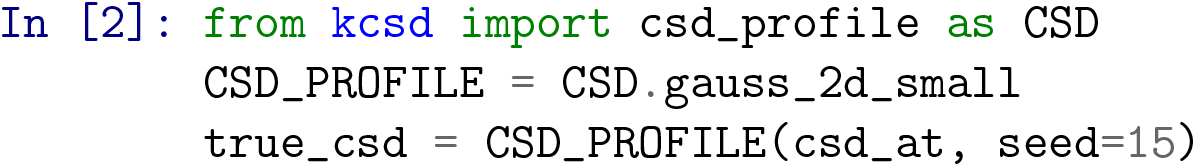

The code below displays this test source as the True CSD. For convenience we define this as a function `make_plot`. The output for this code is shown in Fig. 8A.

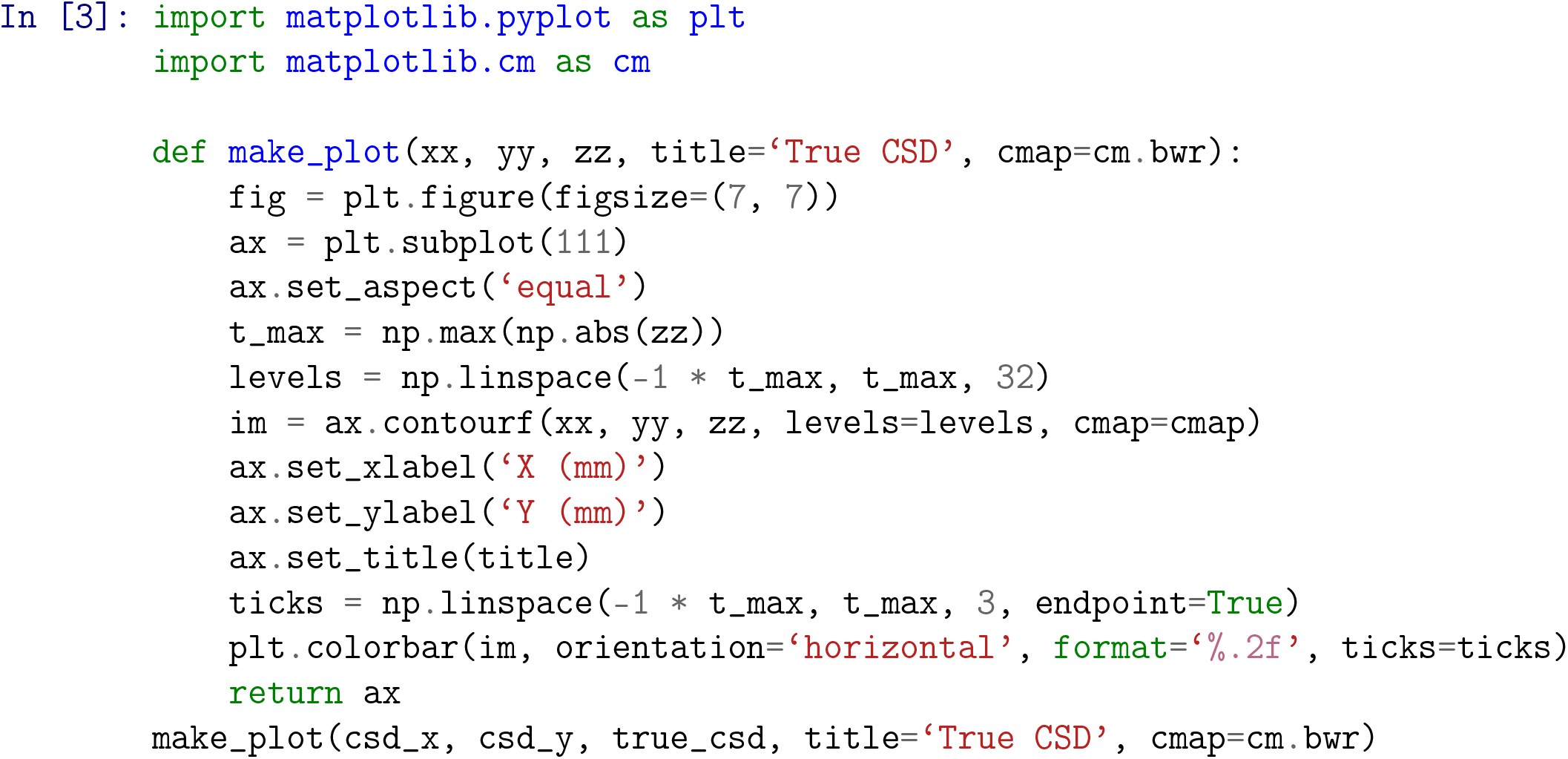

### Place electrodes

We now define the virtual electrodes within the region of interest. We place them between `0.05` *mm* and `0.95` *mm* of the region of interest, with a resolution of `10` (as indicated by `10j` in `mgrid`) in each dimensions, totalling to `100` electrodes. Notice that the electrodes do not span the entire region of interest. Although in this example the electrodes are distributed on a regular grid, this is not required by the kCSD method as it can handle arbitrary distributions of electrodes.

~~~
In [4]: ele_x, ele_y = np.mgrid[0.05: 0.95: 10j,
0.05: 0.95: 10j]
ele_pos = np.vstack((ele_x.flatten(), ele_y.flatten())).T
~~~

### Compute potential

To obtain the potential, `pots`, at the given electrode positions due to the current sources that were placed in the previous steps we use the function `forward_method`. We assume the sources are localized within a slab of tissue of thickness 2`h` on top the MEA (See Łęski, Pettersen, et al., 2011; Ness et al., 2015 and Methods). We also assume infinite homogeneous medium of conductivity `sigma` equal to `1` *S/m*. Finally, we assume that the electrodes are ideal, point-size and noise-free.

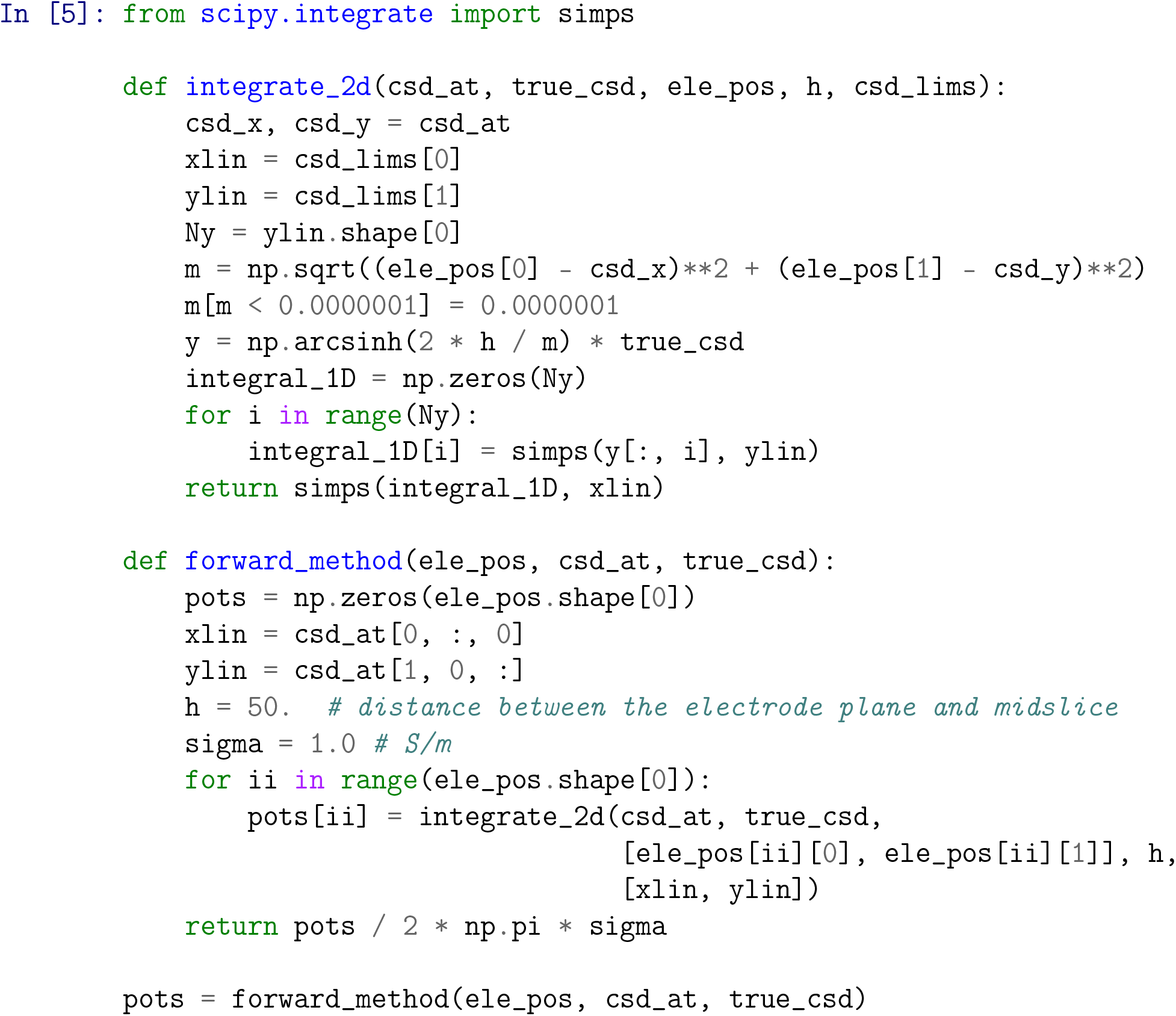

To visualize the potential, we interpolate the hundred values computed at the electrodes positions with interpolate.griddata function. Note that the kCSD estimation uses only the potential recorded at the electrode positions. To distinguish between the potentials and CSD plots we use different colormaps. The electrodes are marked with dots in this plot. The output from this step is shown in Fig. 8B.

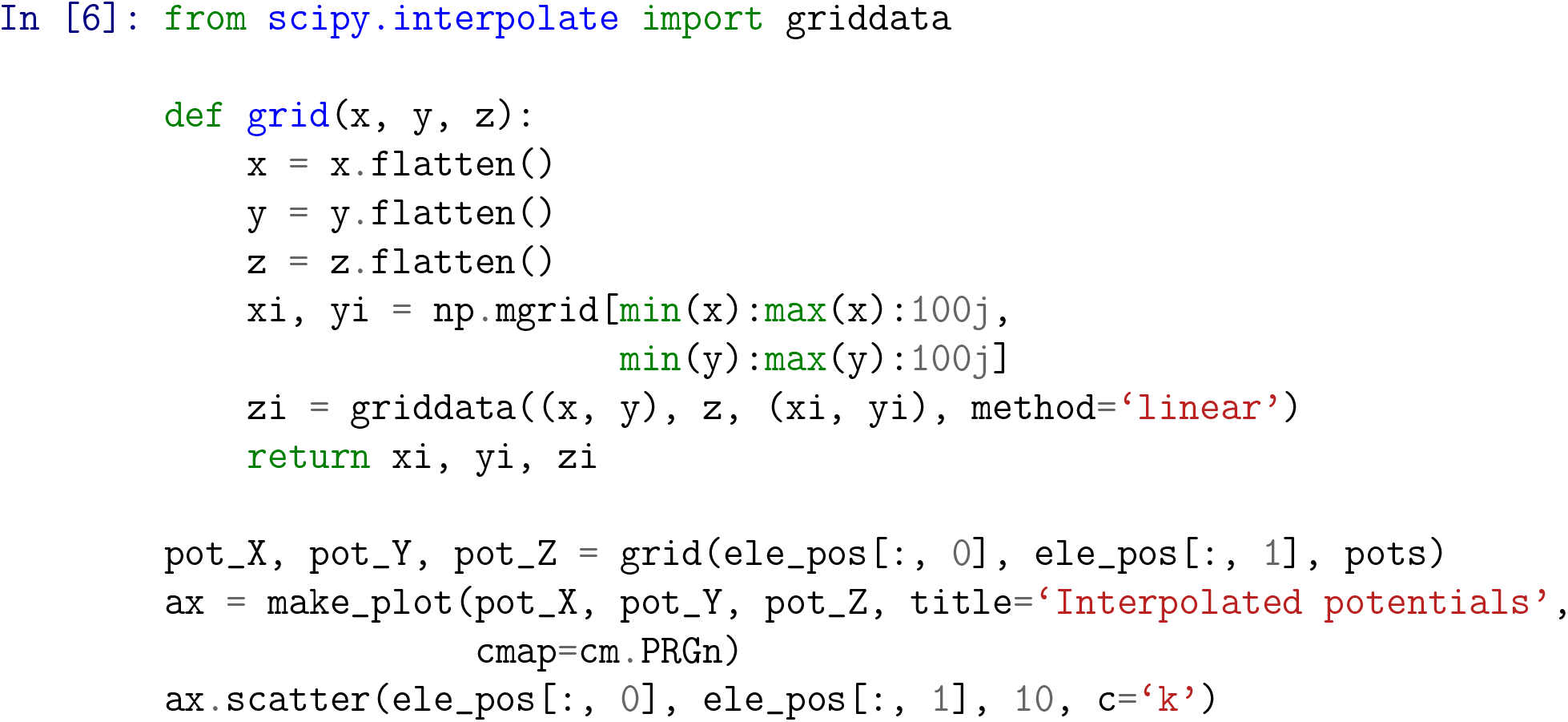

### kCSD method

Here we illustrate the most basic estimation of CSD with the kcsd library. Since our example is two dimensional the relevant method is `KCSD2D`. For convenience we encapsulate the actual method call with parameters being set inside a function `do_kcsd`. We first set `h` and `sigma` parameters of the forward model. Then we restrict the potentials to the first time point of the recording. For typical experimental data the shape of this matrix would be *N*_*ele*_*×N*_*time*_, where *N*_*ele*_ is the number of electrodes and *N*_*time*_ is the total number of recorded time points. Next, we call the `KCSD2D` class with the relevant parameters. The only required parameters are the electrode positions, `ele_pos`, and the potentials they see, `pots`. We can also provide the parameters for the forward model, `h` and `sigma` here. We define a rectangular region of estimation by setting the values `xmin`, `xmax` and `ymin`, `ymax`. The number of basis functions, `n_src_init` is set to `1000`, basis functions are of the type `gauss`, and the width of the Gaussian basis source `R_init` is set to be `1`. Finally, estimated CSD is stored as `est_csd`.

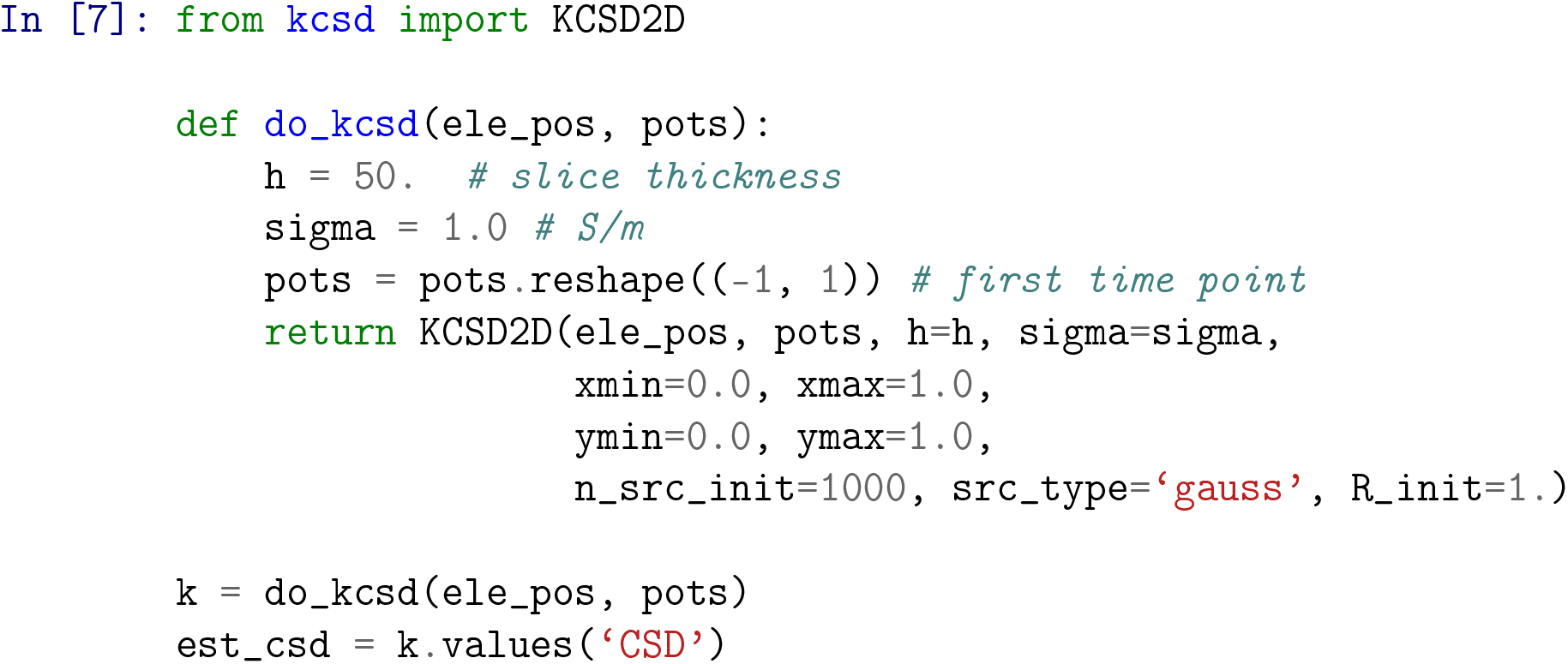

Estimated current sources are shown in Fig. 8C. Compare this to the True CSD obtained before, Fig. 8A. Observe that the estimation is not very faithful. This is caused by the ground truth varying significantly in the scale of a single inter-electrode distance. In the next step we will use cross-validation to select better reconstruction parameters.

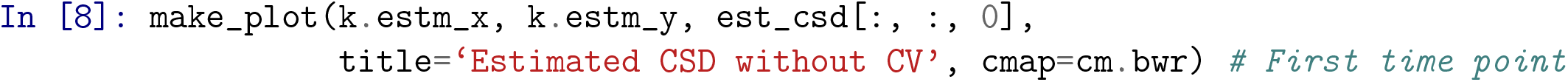

### Cross validation

Leave-one-out cross-validation is performed with a single line command. In this procedure we scan a range of `R` values which set the size of the Gaussian basis functions and the regularization parameter *λ* values. At the end of this step we obtain the optimal parameters that would correct for overfitting. The function outputs the progress of the cross-validation step and displays the optimal candidates in the last line. Alternatively, one could use the L-curve method to find these optimal parameters. Fig. 8D shows the kCSD reconstruction obtained after cross-validation. We find that this estimation of the current sources resembles the True CSD better.

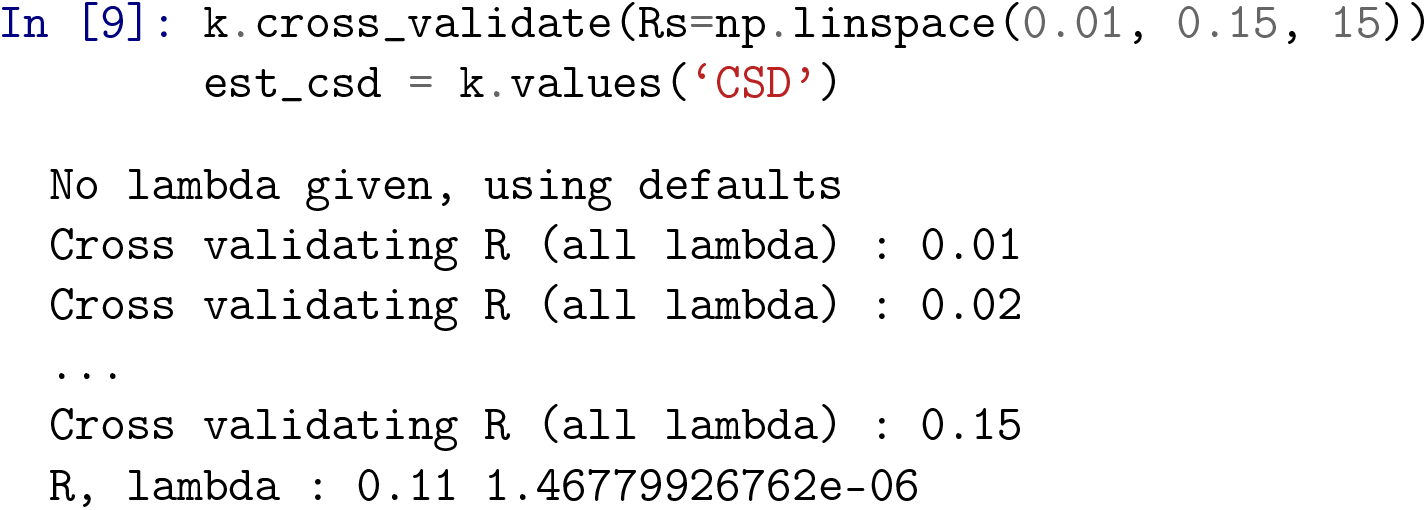

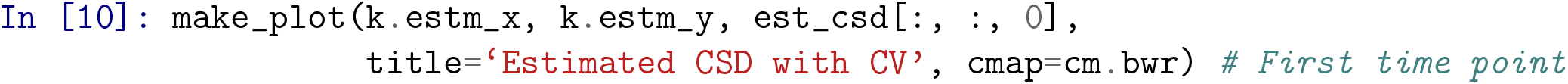

#### Noisy electrodes

Until now we assumed noise-free data, however, experimental data are always noisy. In this section we investigate how noise affects the kCSD estimation. We first show how to compute the reliability map which we introduced before, Eq. (21). Then we discuss reproducible generation of noisy data with varying noise amplitude. Finally, we study the error in the reconstruction as a function of changing noise level.

#### Reconstruction quality measure

To assess the estimation quality we measure the point-wise difference between the true sources and the sources reconstructed with the kcsd. We define a function `point_errors` which takes the `true_csd` and the `estimated_csd` as the inputs, normalizes them individually, and computes the Frobenius norm of their difference.

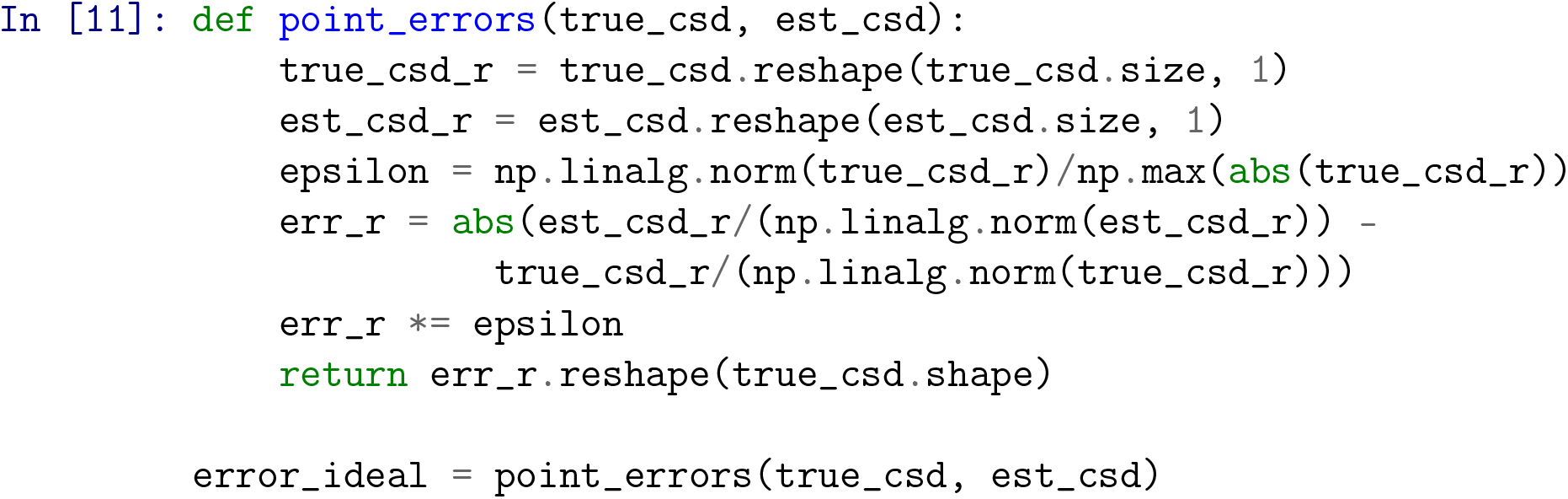

We visualize this difference as before, except we use greyscale colormap to display the intensity of the reconstruction error. For convenience we define the plotting in a function called `make_error_plot`. The output from this step is shown in Fig. 9A.

**Figure 9.**
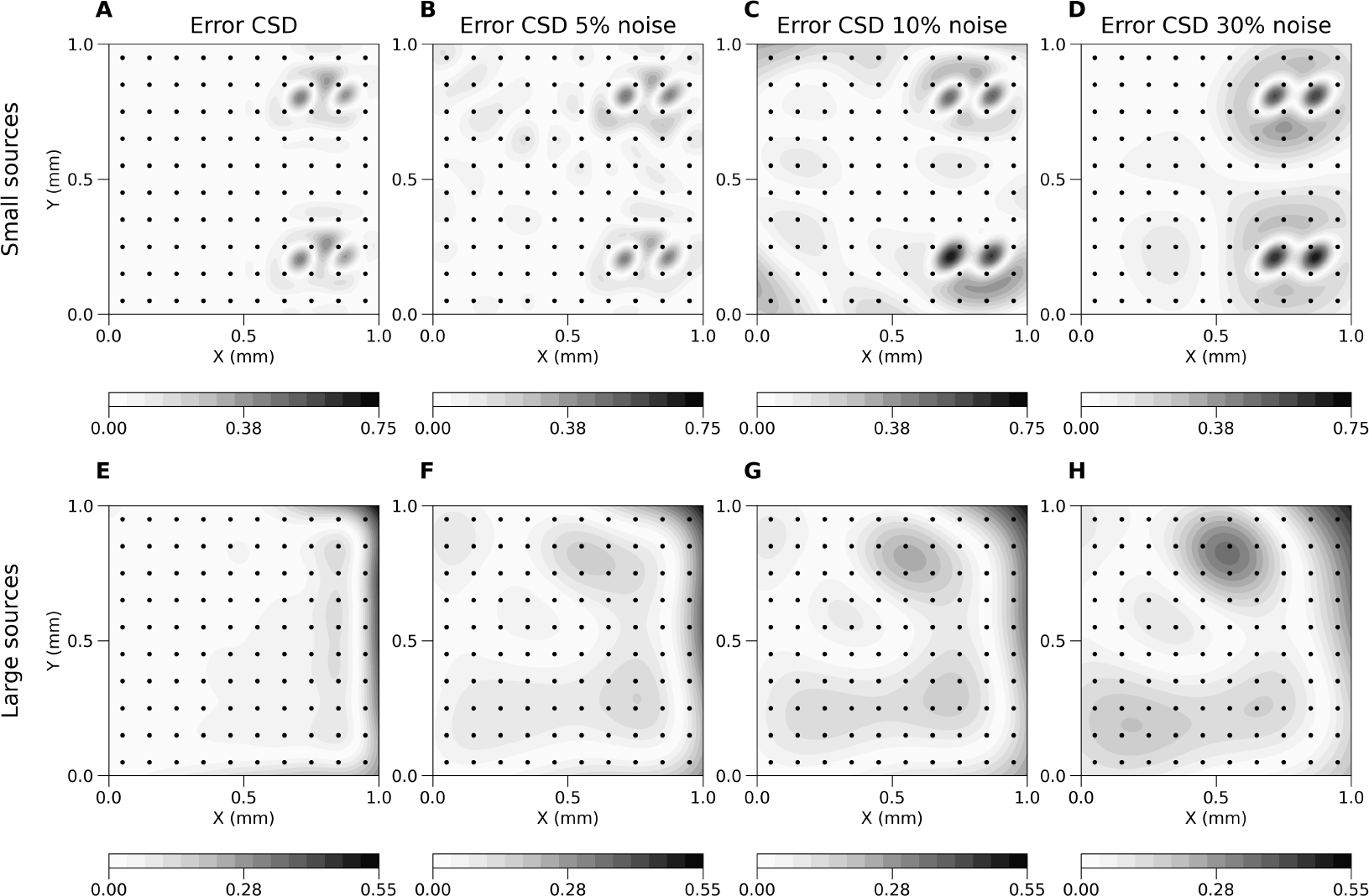
Noisy electrodes. A) The error between the True CSD and the estimation obtained with kCSD for a 2 dimensional small Gaussian current source, using the csd seed of 15. The electrodes in this case are assumed to be noise-free. B, C, D) Same as A, however, noise is added to the recorded potentials, whose magnitude is 5%, 10%, or 30%, respectively. E-H) Analogous to A–D, except in this case large Gaussian sources with seed 6 were used.

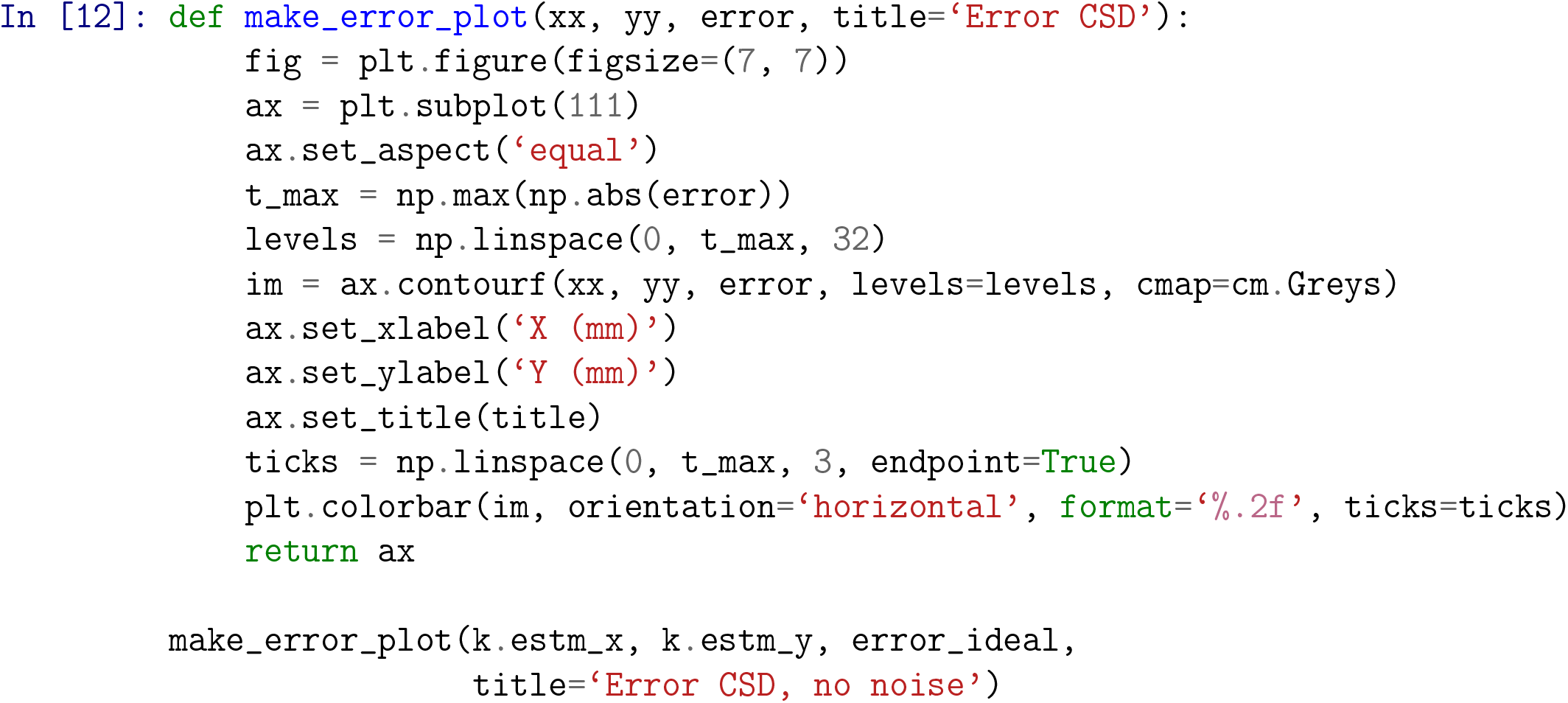

#### Noise definition

To study resilience of the reconstruction against the noise in a controlled way we seed the random number generator in the function `add_noise`. We consider normally distributed noise with the mean and standard deviation set by reference to the recorded potentials.

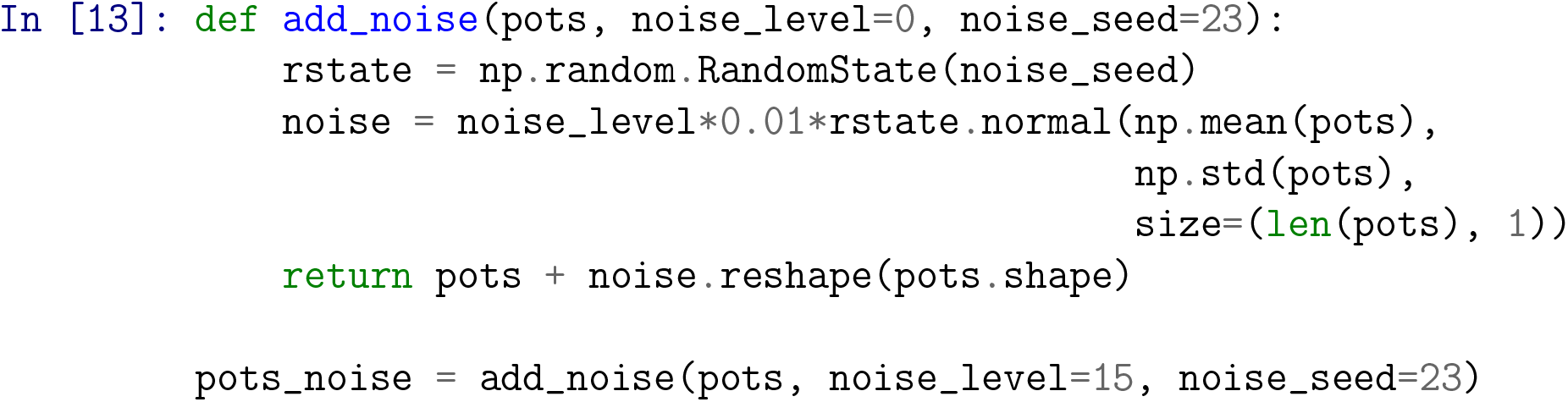

#### Source reconstruction from noisy data

With these tools we can study the effects of noise on the reconstruction. We now generate noise for a given noise level between 0 and 100, add it to the simulated potential, and estimate CSD from these noisy potentials. We can then use the error plots to compare the reconstruction with the True CSD. Notice that the parameters giving best reconstruction obtained for noisy data in general will be different from those obtained for clean potentials to compensate for noise.

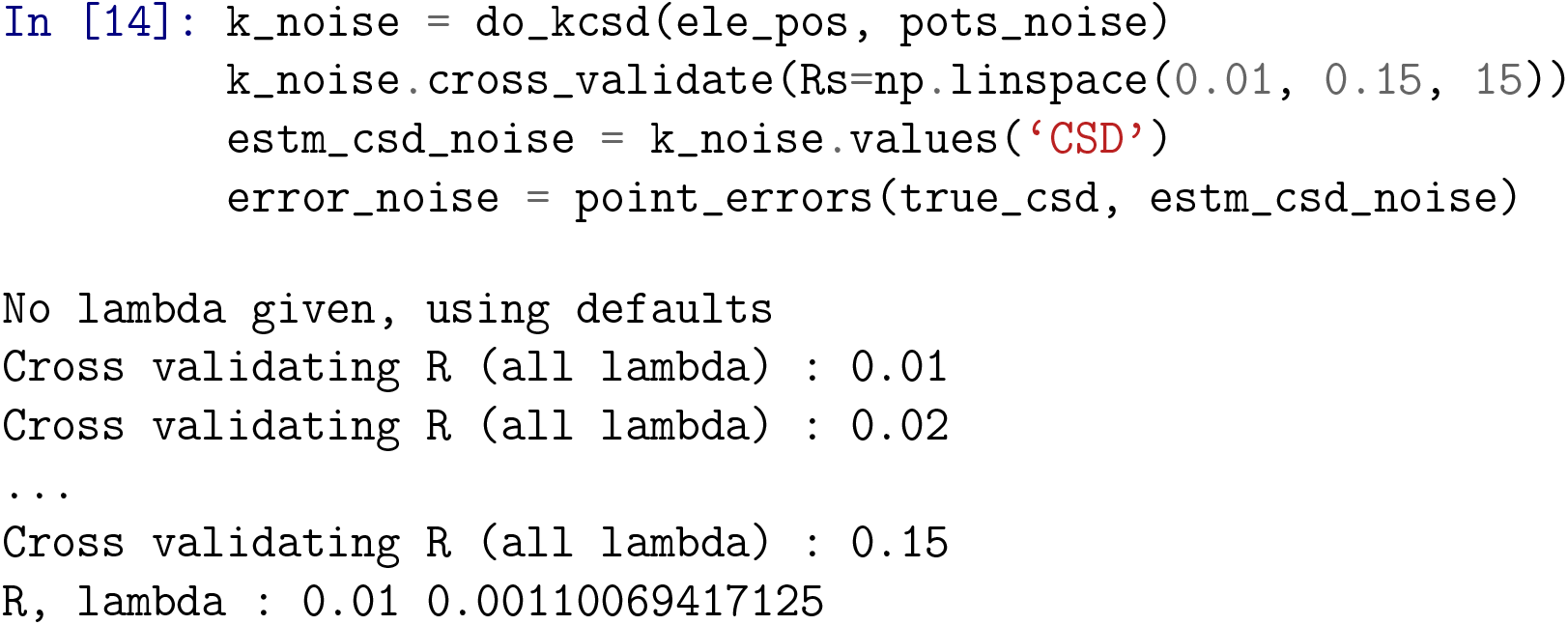

We can display this error with the `make_error_plot` plotting function which we defined earlier. Changing the `noise_level` and the `noise_seed` affects the reconstruction, but the error depends also on the sources, so changing the True CSD type to a `gauss_2d_large` or changing `csd_seed` will lead to different results. This is illustrated in Fig. 9A–D for small Gaussian sources, and Fig. 9E–H for large Gaussian sources, with varying noise levels. The actual ground truth and reconstructions are shown in Fig. 8.

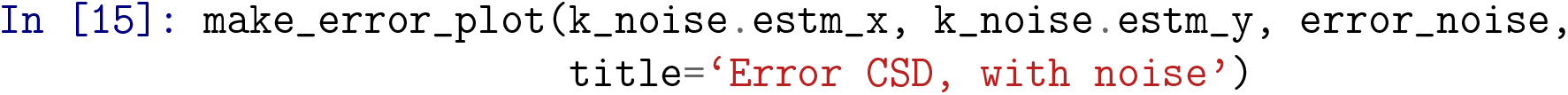

### Broken electrodes

It is often the case that due to experimental constraints, some subset of recordings are discarded in the final analysis. This can happen when some electrodes are used for stimulation and cannot be used for recording, or due to bandwidth limitations requiring a compromise between sampling rates and the number of simultaneously recording electrodes, or in the event of an electrode break down. In this section of the tutorial we discuss how to handle such cases and to estimate the errors in reconstruction despite the loss of recordings. We first show how we remove recordings from selected (broken) electrodes from considered data. Then we calculate the estimation error for a given source for data from such a damaged setup. Finally, we compute the average error across many sources from such an incomplete setup. Note that kCSD reconstruction is designed to work with arbitrary electrode setups and removing specific electrodes does not change the situation significantly. We focus on broken electrodes as it is a common situation in practice and deserves consideration. This may be used to gain intuition regarding ways in which `CSD` reconstruction may go wrong, due to slight disturbances in a familiar setup.

#### Remove broken electrodes

To test the effects of removed electrodes on reconstruction from a given setup we simulate this with a function `remove_electrodes` that takes all the electrode positions for this setup and the number of electrodes that are to be removed. In this example we remove the electrodes randomly. Like we did previously, to facilitate repeatability we also pass a `broken_seed` variable, so that at each subsequent run the same electrodes are discarded. By changing this seed we select a different set of electrodes for removal.

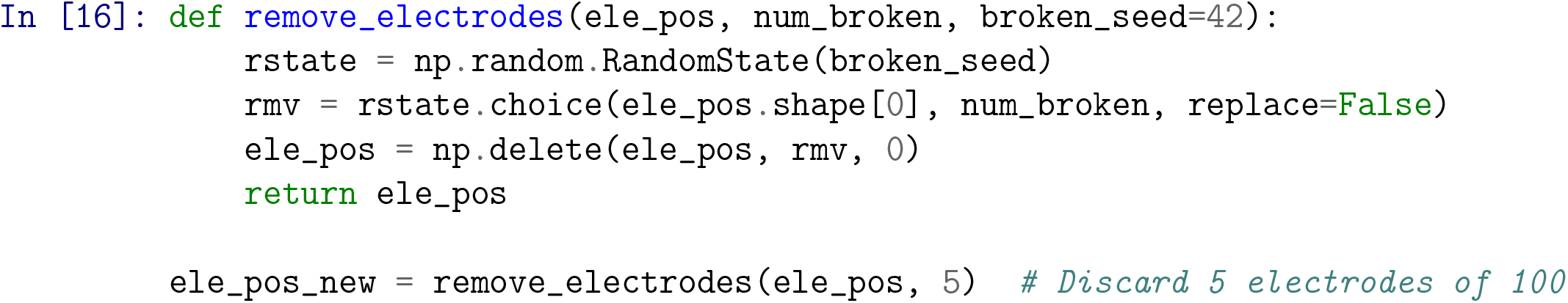

#### Error in estimation with broken electrodes

After removing the broken electrodes we compute the estimation error to gauge the effect of electrode removal on reconstruction. Here, a fuction `calculate_error` takes a `csd_seed` as an input, which selects a specific ground truth source, and all the remaining electrode positions, `ele_pos`. The function computes the True CSD for a `gauss_2d_small` type source, computes the potential at these electrode locations, performes kcsd estimation from these data, and computes the error in the estimation of the true csd.

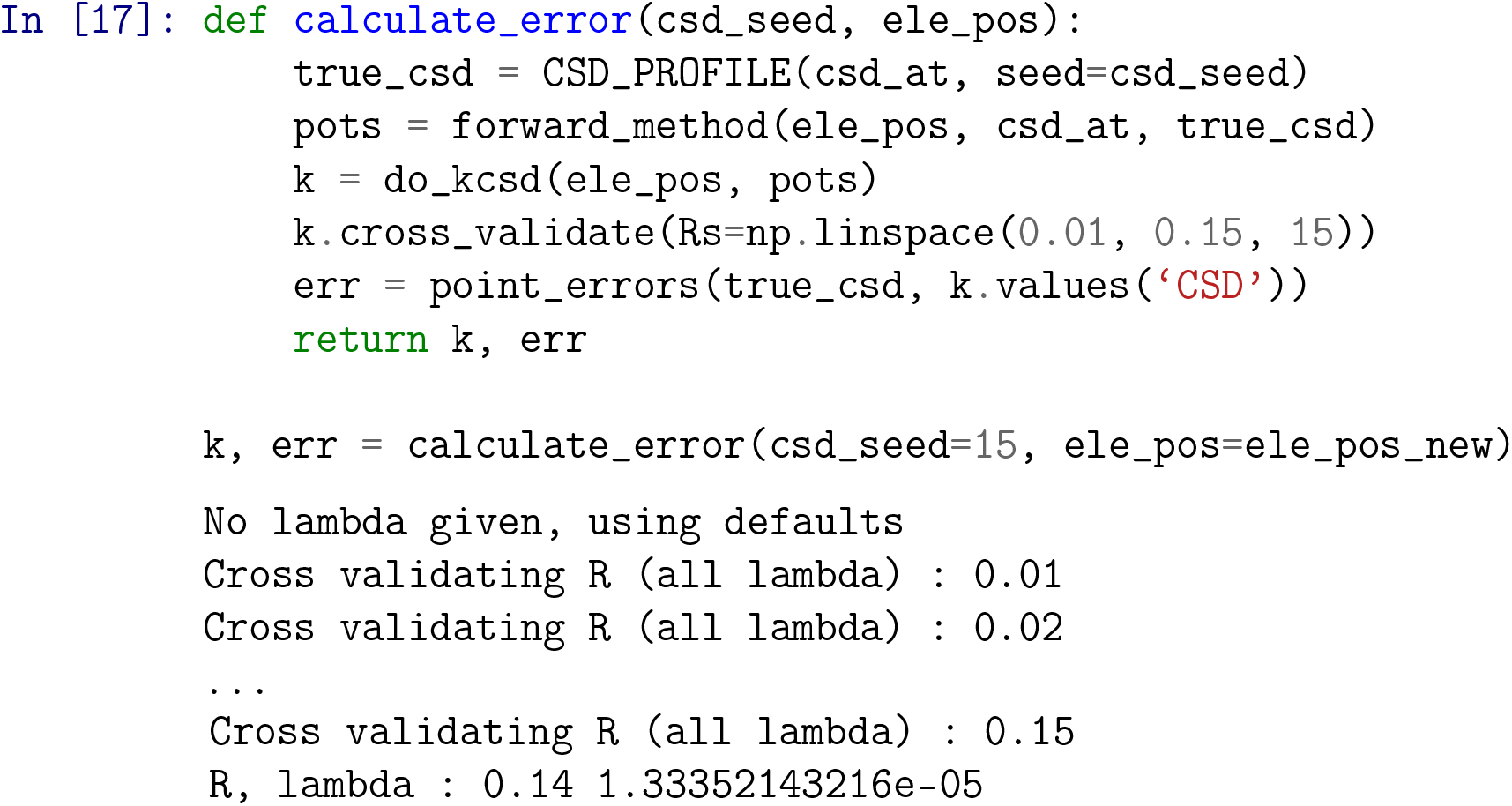

Below (Fig. 10) we plot these errors. We also display the electrodes which were used in the kcsd estimation.

**Figure 10.**
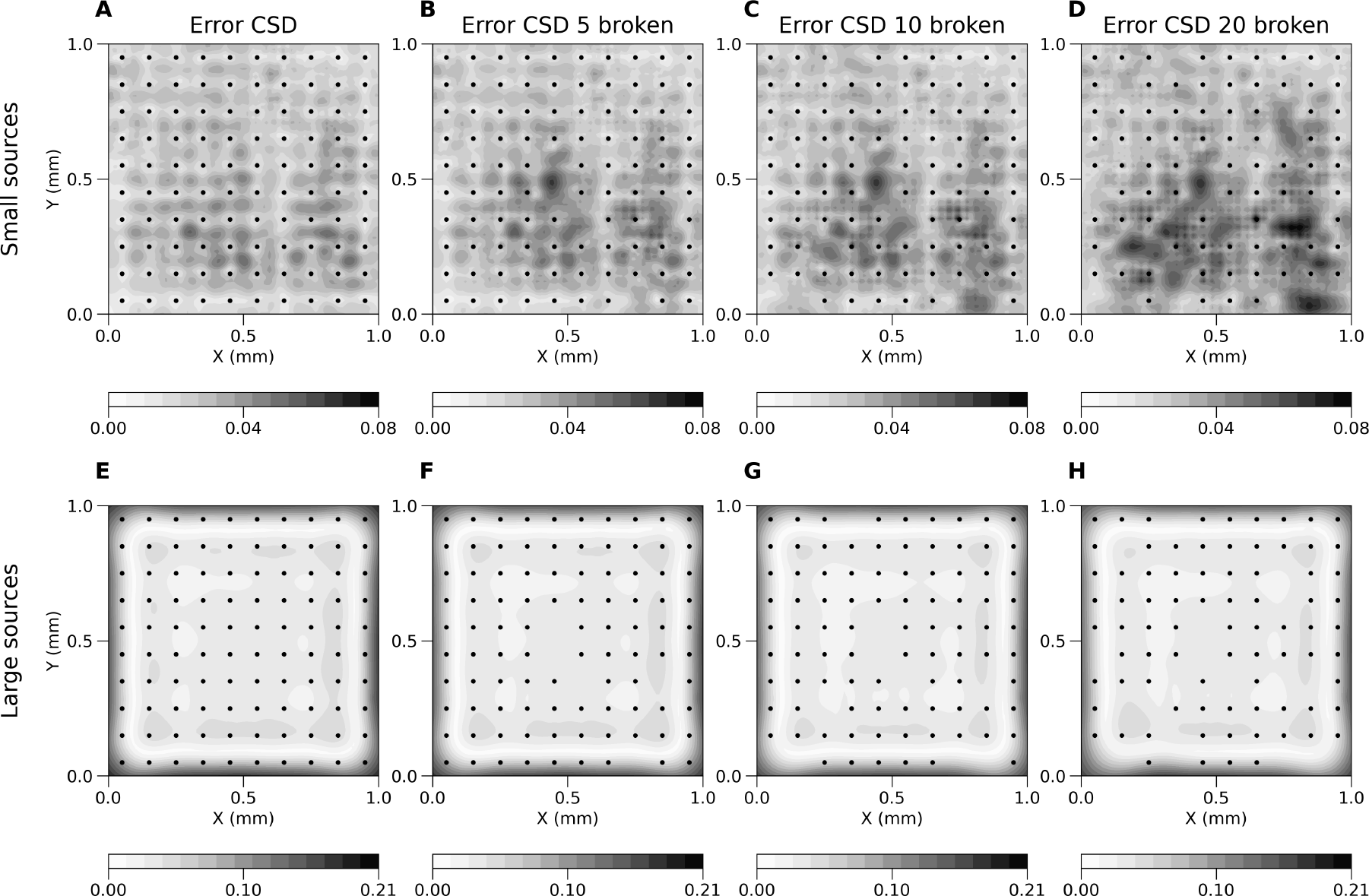
Broken electrodes. A) Shows the average error between the True CSD and the CSD estimated with kcsd for 100 random small Gaussian current sources. B) The same average error as in A, except in this case 5 electrodes were discarded in the estimation. Likewise for C and D, where 10 and 20 electrodes out of the 100 were considered broken. E-H) analogous to A–D, except in this case we show the averages for 100 large Gaussian current sources.

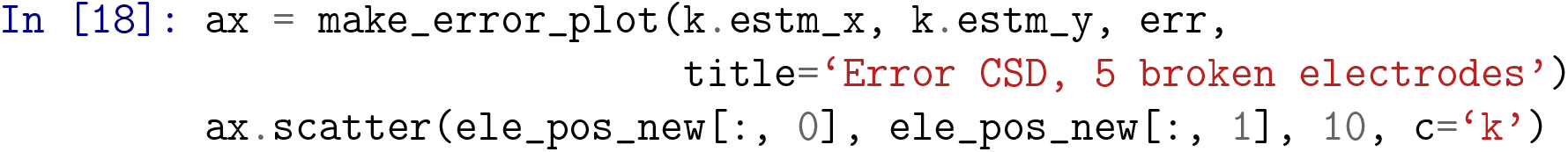

#### Average error for multiple sources

As we can see, the estimation error depends on the test current sources used. To better understand the effects of the setup we compute the average error across multiple sources. As an example here we show this for two seeds. In principle, any type and number of sources may be tested, as we showed before in analysis of reliability maps. This step is computationally expensive, however, it would normally be carried out only once for a given electrode design configuration. We believe this approach offers useful diagnostics and builds intuition regarding the estimation power for the given setup.

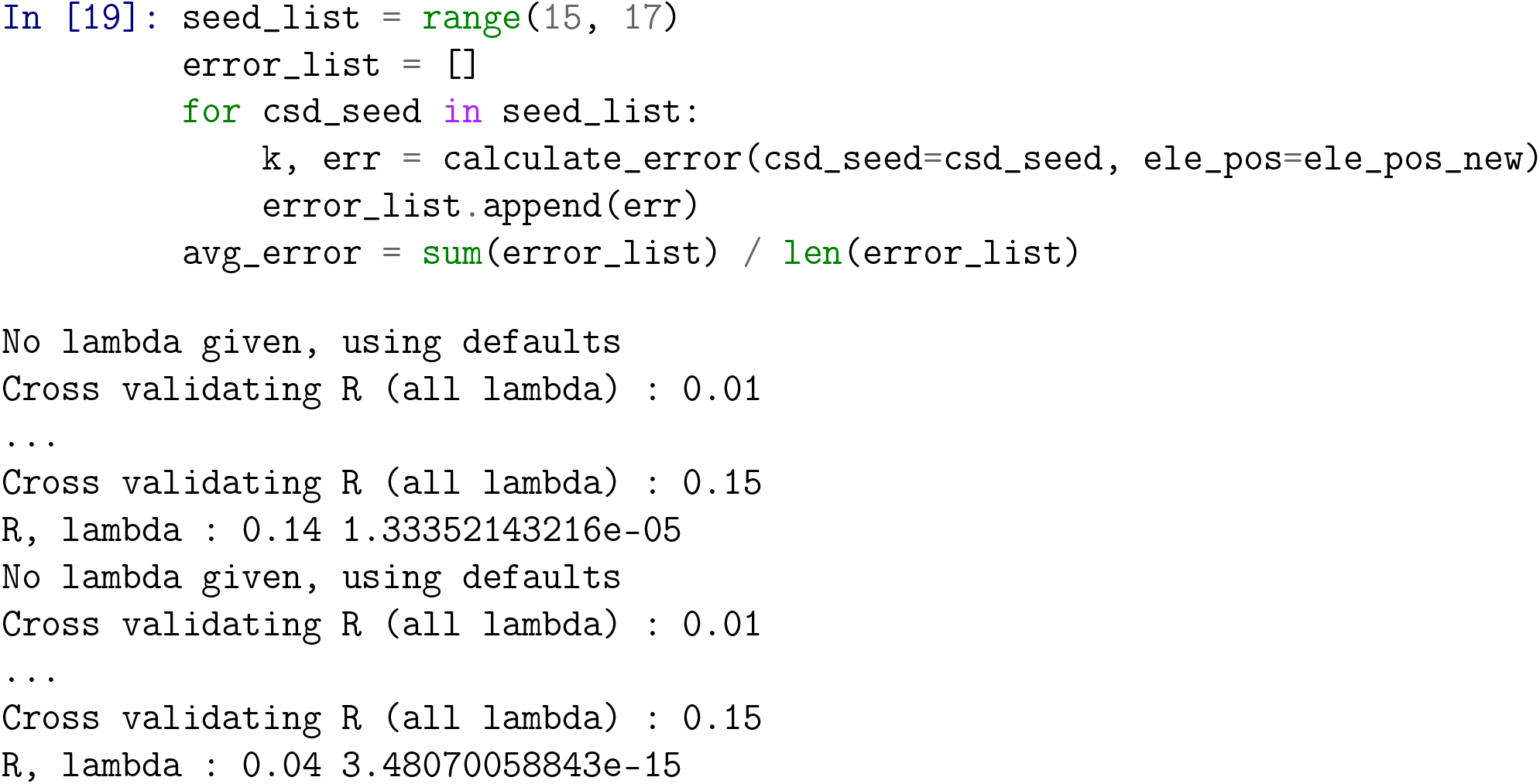

In Fig. 10A–D we show this for the case of 0, 5, 10 and 20 broken electrodes, when the average error for 100 small Gaussian sources was considered. In Fig. 10E–H we show the same for large Gaussian sources.

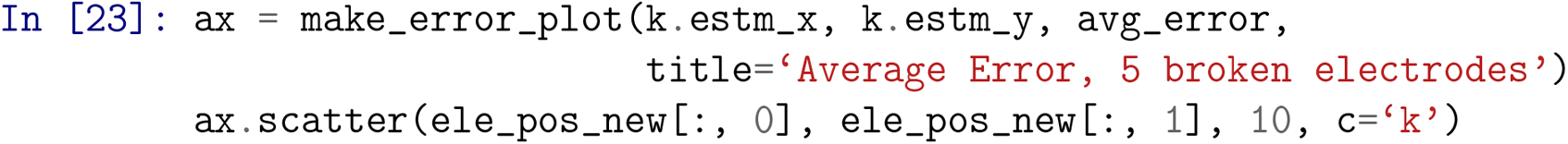

### Supplementary Materials

#### S1 Fig

##### Error propagation maps in a 1D example

**Figure 11.**
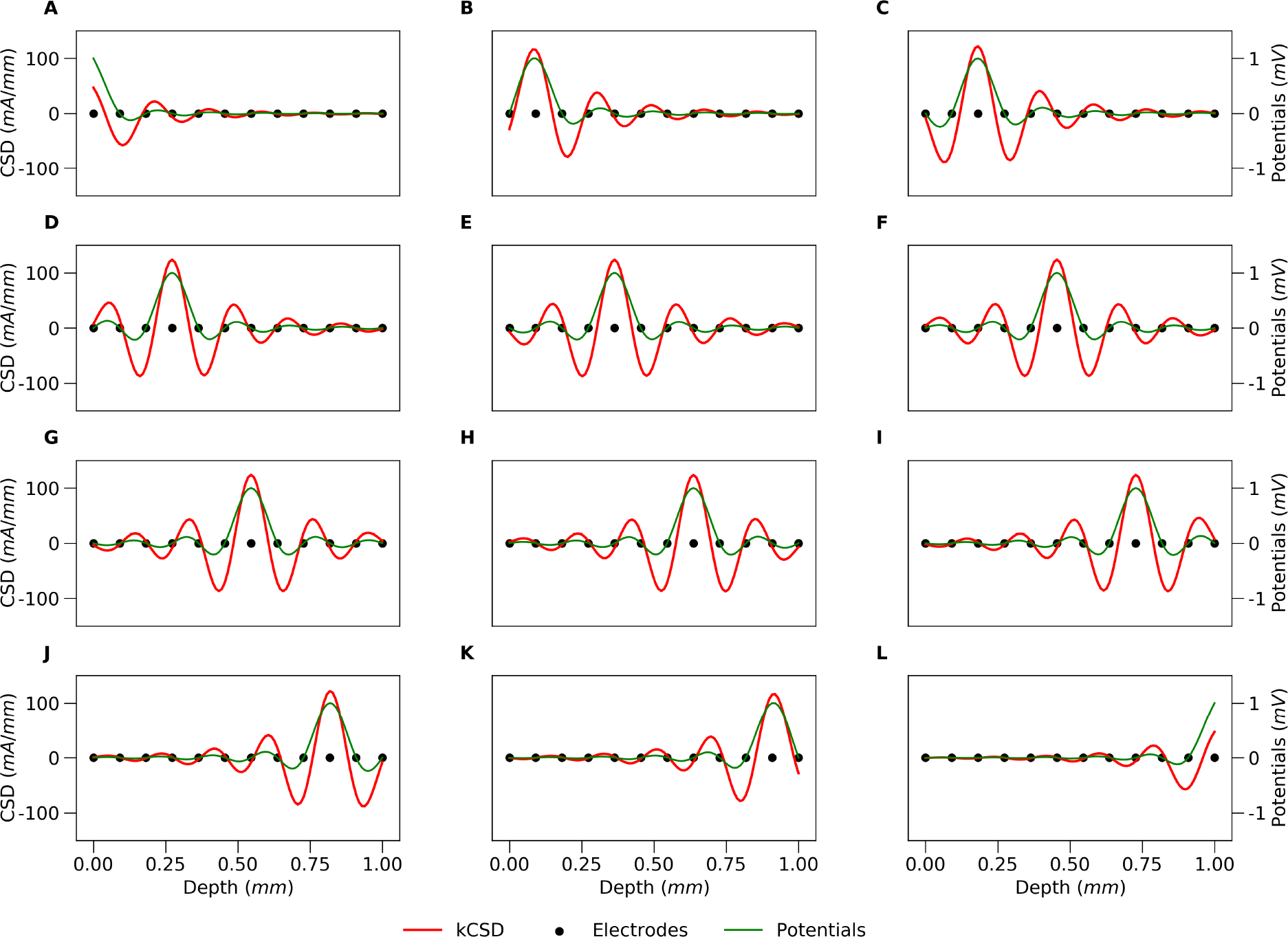
**Error propagation maps** for 1D regular grid of 12 electrodes Every panel represents the CSD contribution (red line) of the potential measured at the corresponding electrode, for which the potential is 1 (green line).

#### S2 Fig

##### An example of 3D source reconstruction

**Figure 12.**
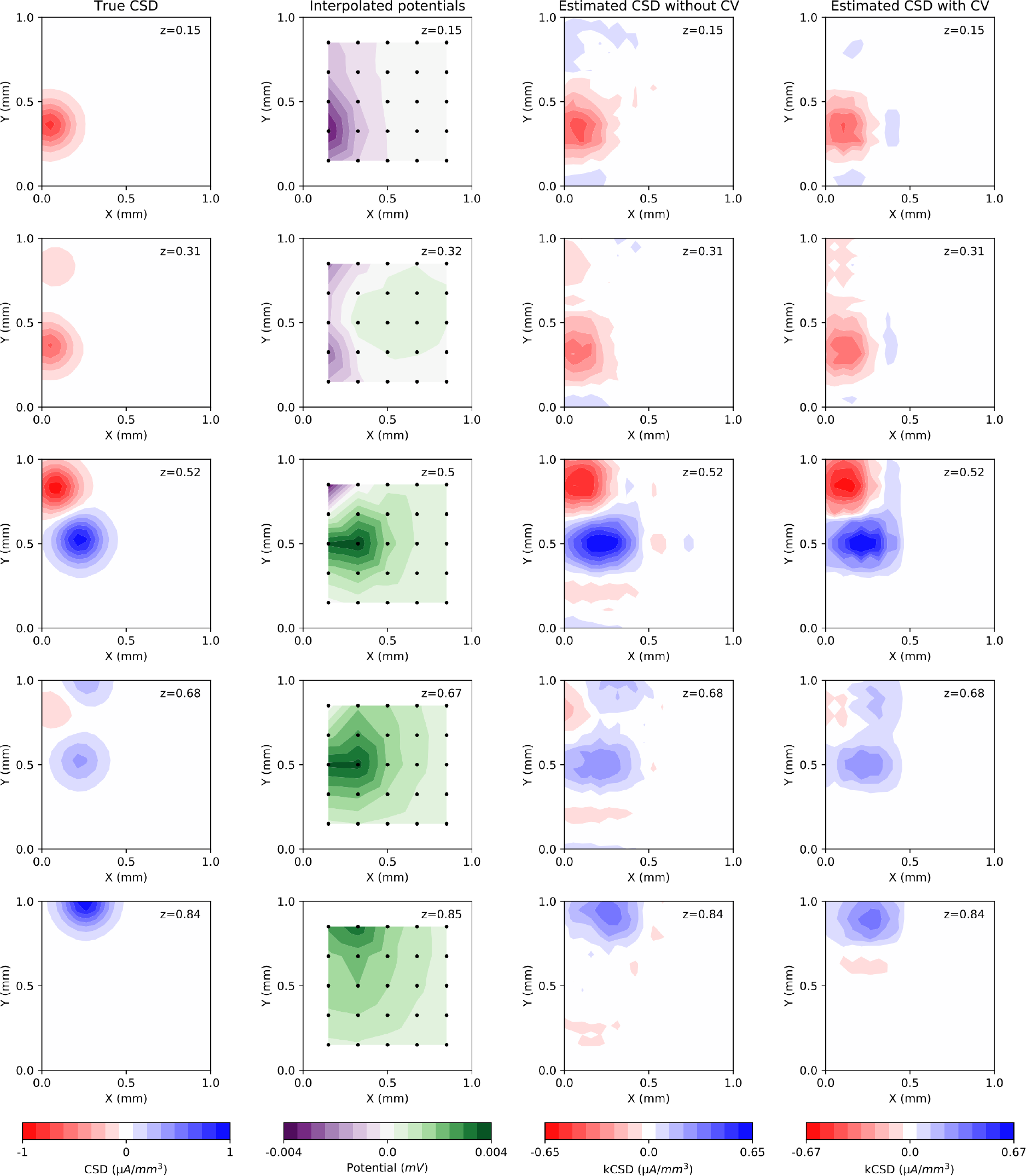
**An example of 3D kCSD source reconstruction**. Each column shows five consecutive parallel cuts through a box of size 1. A) Ground truth for the CSD seed of 16. B) Estimated potential; black dots indicate electrodes where potential is collected for further reconstruction. C) 3D kCSD reconstruction from the measured potentials, *λ* = 0. D) 3D kCSD reconstruction with cross-validation.

#### S3 Fig

##### An example of skCSD source reconstruction

**Figure 13.**
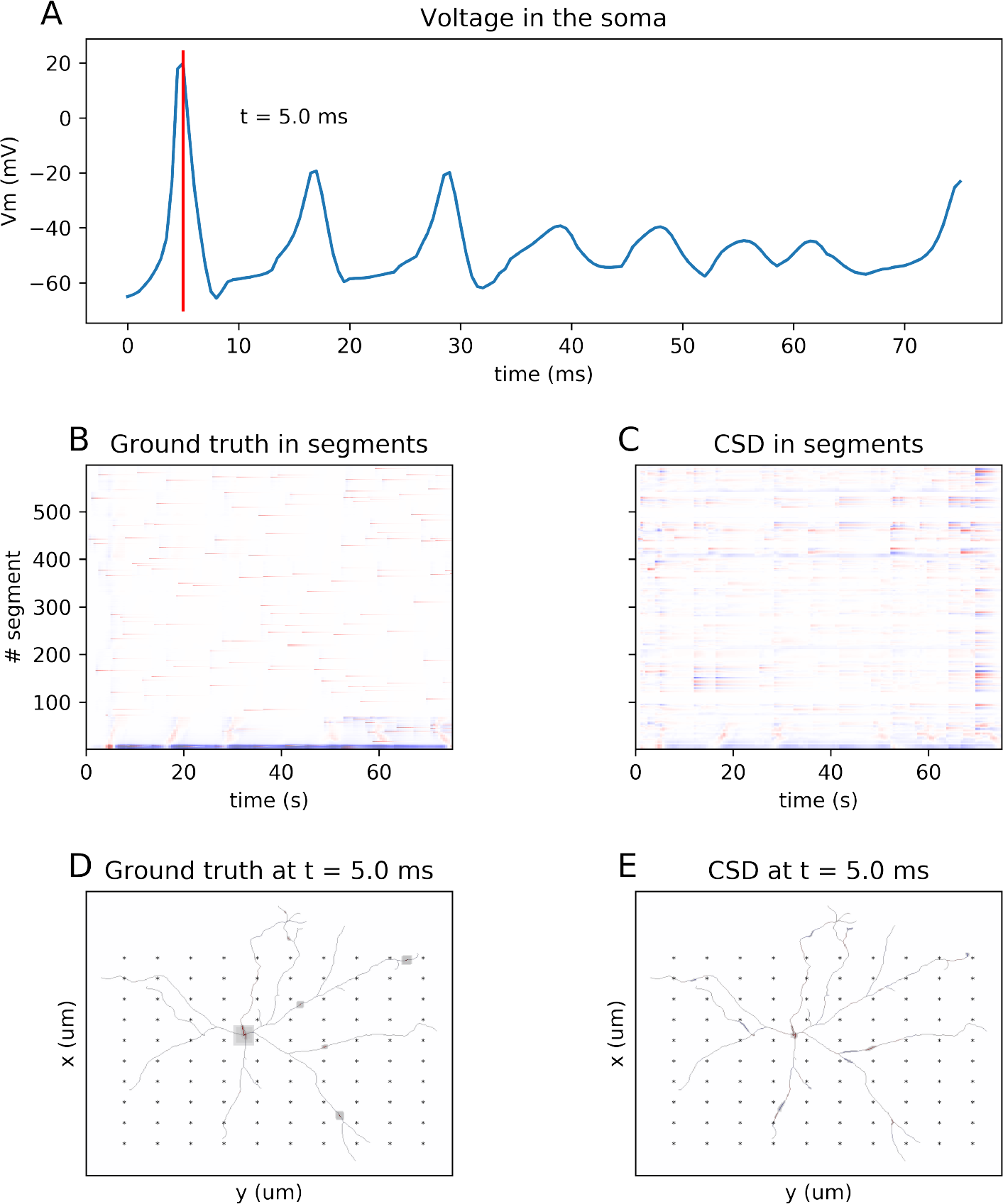
**An example of skCSD reconstruction** Cserpan et al., 2017 of somatic current injection together with random synaptic input patterns for a retinal ganglion cell model. A) Somatic membrane potential. B) Current density and its C) skCSD reconstruction in the segment space. Projection of D) ground truth and E) skCSD reconstruction on the neuron’s morphology at 5 s of the simulation. We simulated a multicompartmental model of a mouse retinal ganglion cell (morphology Kong et al., 2005 obtained from NeuroMorpho.Org Ascoli, 2006) with Hodgkin-Huxley sodium, potassium, and leakage channels in the soma (hh mechanism) in NEURON simulation environment. For calculation of the measured extracellular potentials we used LFPy package Lindén, Hagen, et al., 2013. The model neuron was stimulated by an injection of oscillatory current to the soma (with frequency of 24.5 1/ms and amplitude of 3.6 nA) together with random synaptic inputs (weight of 0.04 *μS*) to the dendritic tree. The activity of the model neuron was measured by a rectangular grid of 100 electrodes (10*×*10, -400*×μm* 400*μm*). The figure corresponds to Fig. 8 from (Cserpan et al., 2017).

## Footnotes

1 supports also Python 2.7 version

2 In the special case when *ε*_*i*_ are mutually independent and of equal variance *σ*^2^, the map of CSD measurement uncertainty can be calculated as a diagonal of Cov[**C**^*\**^] = **EE**^*T*^ *σ*^2^.

